# Cell type- and region-specific modulation of cocaine seeking by micro-RNA-1 in striatal projection neurons

**DOI:** 10.1101/2021.05.27.445939

**Authors:** Benoit Forget, Elena Martin Garcia, Arthur Godino, Laura Domingo Rodriguez, Vincent Kappes, Pierre Poirier, Andry Andrianarivelo, Eric Senabre Marchan, Marie-Charlotte Allichon, Mélanie Marias, Peter Vanhoutte, Jean-Antoine Girault, Rafael Maldonado, Jocelyne Caboche

**Author notes:** equal contribution (Benoit Forget, Jocelyne Caboche).

## Abstract

The persistent and experience-dependent nature of drug addiction may result in part from epigenetic alterations, including non-coding micro-RNAs (miRNAs), which are both critical for neuronal function and modulated by cocaine in the striatum. Two major striatal cell populations, the striato-nigral and striato-pallidal projection neurons, express respectively the D1 (D1-SPNs) and D2 (D2-SPNs) dopamine receptor, and display distinct but complementary functions in drug-evoked responses. However, a cell-type-specific role for miRNAs action has yet to be clarified. Here, we evaluated the expression of a subset of miRNAs proposed to modulate cocaine effects in the nucleus accumbens (NAc) and dorsal striatum (DS) upon sustained cocaine exposure in mice and showed that these selected miRNAs were preferentially up-regulated in the NAc. We then focused on miR-1 considering the important role of some of its predicted mRNA targets, such as *fosb* and *npas4*, in the effects of cocaine. We validated these targets *in vitro* and *in vivo* and further showed that overexpression of miR-1 in D1-SPNs of the DS reduced cocaine-induced CPP reinstatement, whereas it increased cue-induced reinstatement of cocaine-SA, without affecting other cocaine-mediated adaptive behavior. In addition, miR-1 overexpression in D2-SPNs of the DS reduced the motivation to self-administer cocaine but did not modify other measured behaviors. Together, our results highlight a precise cell-type- and region-specific control of relapse to cocaine-seeking behaviors by miR-1, and illustrate the importance of cell-specific investigations.

## INTRODUCTION

Drug addiction is a clinically devastating neuropsychiatric disorder thought to result from neural adaptations at the molecular, cellular, and circuit levels, following repeated drug exposure [1]. Drugs of abuse initially affect neuronal plasticity in the mesocorticolimbic system corresponding to dopamine (DA) neurons of the ventral tegmental area (VTA) projecting onto the ventral part of the striatum, the *nucleus accumbens* (NAc), and the prefrontal cortex [2,3]. The striatum plays crucial roles in diverse behavioral processes including movement control and reward-dependent learning [4–6]. While the NAc underlies the primary reinforcing effects of drugs of abuse such as cocaine, the emergence of compulsive cocaine seeking habits, a hallmark of drug addiction, is associated with a shift to the dorsal striatal (DS) control over behavior [7,8].

In the DS, and to a lesser extent in the NAc, striatal projection neurons (SPNs) are comprised of two segregated GABAergic neuronal populations: the SPNs expressing the DA-D1 receptor subtype (D1- SPNs) that project directly to the internal segment of the *globus pallidus* (GPi) and the *substantia nigra pars reticulata* (SNr), and the SPNs expressing the DA-D2 receptor subtype (D2-SPNs), which project mainly to the external segment of the *globus pallidus* (GPe). Furthermore, these two subpopulations exert distinct, albeit complementary, functions. In the DS the D1-SPNs represent the “direct pathway”, which facilitates the initiation and execution of voluntary movement, while the D2- SPNs of the “indirect pathway” inhibit movement [9,10]. However, both pathways are necessary for action initiation [11]. The direct and indirect pathways are also thought to work in opposing manners in reward-based learning, with the direct pathway mediating reinforcement and the indirect pathway mediating punishment and aversion [5,12], although such a dichotomy has been recently challenged [13–15]. Several studies investigated the participation of D1- and D2-SPNs in behaviors related to cocaine exposure [16]. For example, activation of D1-SPNs in the NAc increases the formation of cocaine reward– context associations, while activation of D2-SPNs in the same structure decreases cocaine reward and self-administration [17,18]. D1-SPNs in the NAc encode information related to the association between cocaine and contextual cues and drive cocaine seeking induced by those cues [19]. While most of the studies investigating the role of D1- and D2-SPNs in the behavioral effects of cocaine focused on the NAc, a recent work showed a very specific role of D1-SPNs of the dorsomedial striatum in cocaine responses, where their inhibition had no impact on loss of control over cocaine intake, motivation to obtain cocaine or compulsive cocaine self-administration, but specifically reduced cue-induced cocaine-seeking in high-risk addiction groups of rats [20].

The persistent and experience-dependent aspects of the addicted phenotype have suggested a role for epigenetic modifications in drug-induced gene expression [21] as epigenetic mechanisms can register and durably maintain transcriptional states [22]. Among various epigenetic processes, attention has recently been focused on miRNAs, a subcategory of 21-25-nucleotides non-coding RNAs that modulate gene expression by binding to complementary sequences in the 3’ untranslated regions (3’UTR) of many, up to hundreds, target mRNA transcripts per miRNA, thereby inducing mRNA destabilization and subsequent degradation and/or repression of mRNA translation [23]. Changes of miRNAs expression levels have been shown to play a role in multiple neuronal functions and are altered in several psychiatric conditions [24], including addiction [25]. Several studies have provided evidence that miRNA-mediated gene regulation plays an important role in cocaine-induced changes in neurotransmission and behavior [26], but there is so far, no direct evidence regarding cell-type-specificity of cocaine-induced miRNA in the striatum, and more specifically in D1-SPNs versus D2- SPNs. One study began to broadly address this aspect by specific ablation in D2-SPNs of Ago2 [27], a key protein in the processing and production of miRNAs [28–30]. These mice showed loss of motivation to self-administer cocaine, along with a decrease of around one third of miRNAs induced by acute cocaine, thus indicating that miRNA induction occurs in both D1- and D2-SPNs. However, the question of whether and how miRNAs regulation in D1-SPNs versus D2-SPNs controls gene expression and behaviors induced by cocaine, remains to be addressed.

Here we aimed to identify the regional pattern, i.e. DS *versus* NAc, of expression of a set of selected miRNAs, proposed to play a key role in cocaine action or neuronal plasticity after sub-chronic cocaine administrations [24,25,31]. We next focused on miR-1, based on its predicted mRNA targets, including *fosb* and *npas4* mRNAs. We used a validated viral strategy [32–34] in order to thoroughly evaluate miR-1 cell-type-specific roles in D1- or D2-SPNs of the NAc or DS in multiple behaviors related to cocaine addiction. We found a key role of miR-1 overexpression in D1-SPNs of the DS on cocaine-and cue-induced relapse to cocaine seeking, with a reduction of cocaine-induced reinstatement of cocaine seeking in conditioned place preference (CPP) paradigm, and an increase in cue-induced reinstatement of cocaine self-administration (SA). We also observed a reduction of motivation for cocaine self-administration after miR-1 overexpression in D2-SPNs of the DS. Altogether our data unveil a complex cell-type- and region-specific regulatory role for miR-1 in different relapse behaviors, and the importance of cell-specific investigations.

## MATERIALS AND METHODS

More details on the materials and methods used are given in Supplementary Methods.

### Animals and drug treatments

All animal treatments and procedures were conducted in accordance with local and European directives for the care and use of laboratory animals (86/609/EEC) and approved by the institutional French Animal Care Committee (agreement N°2010/63) and by the Spanish local ethical committee (*Comitè Ètic d’Experimentació Animal-Parc de Recerca Biomèdica de Barcelona*, CEEA-PRBB).

### Viral vectors

For the overexpression of miR-1 in D1-SPNs or D2-SPNs of the striatum, we used a combination of AAVs (AAV9-*PPTA*-Cre or AAV9-*PPE*-Cre, respectively; titers: 1 × 10^12^ in combination with AAV9-miR-1^flox^ or AAV9-miR-Scr^flox^ titers: 1 × 10^12^ genomic copies/mL). AAV-injected mice were used for experiments after at least three weeks for the full expression of viral-mediated miRNA expression.

### Stereotaxic injections

Mice were anesthetized using ketamine and xylazine (5:1 in amount; 0.1 mL/10g), placed in a stereotaxic frame (David Kopf, Tujunga, CA), and 1 μl in the NAc or 2 μl in the DS was injected at a constant rate of 0.20 μl/min by using a microinfusion pump (Harvard Apparatus, Holliston, MA). We used the following coordinates to perform bilateral injections according to the mouse brain atlas of Paxinos and Franklin: NAc coordinates: AP +1.4 mm; ML ±0.7 mm; DV −4.4 mm; DS coordinates: AP +0.98 mm; ML ±1.75 mm; DV −3.5 mm.

### Quantitative PCR analysis (qPCR) of miRNA and mRNA levels

Punches of NAc and DS were performed from frozen tissues and total RNA was isolated using miRNeasy MiniKit (Qiagen) according to the manufacturer’s instructions. For mRNA levels analysis, total RNA (100 ng) isolated from each brain sample was reverse-transcribed with oligodT primers using RevertAid First Strand cDNA Synthesis Kit (ThermoFisher). For miRNA levels analysis, total RNA (100 ng) was reverse-transcribed using the same kit with a combination of specific RT-primers for each miRNA of interest (0.5 pmol of each RT-primer), as previously described [35], allowing high specificity, sensitivity and homogeneity. qPCR was performed with a LightCycler 480 device (Roche) using LightCycler SYBR Green I Master Mix (Roche) and specific couples of primers; samples were run in triplicates. The primers sequences for miRNAs and mRNAs are presented in supplementary **Figure 1a and b**, respectively.

### Luciferase assay to assess the interaction between miR-1 and the 3’UTR of *Bdnf*, *FosB* and *Npas4* in HEK cells

The pmirGLO-3’UTR_*Bdnf*, pmirGLO-3’UTR_*FosB* and pmirGLO-3’UTR_*Npas4* plasmids were purchased from Creative Biogene and the mimick-miR-1a-3p and mimick-miR-negative control from Qiagen (miScript miRNA Mimics). The protocol used is described in DOI: 10.21769/BioProtoc.420. The mean ratio of luminescence from the experimental reporter (firefly) to luminescence from the control reporter (Renilla) was calculated for each triplicate and normalized to the ratio of control wells (miRNA negative control which does not target the UTR).

### Cocaine-induced locomotor sensitization

The locomotor activity was evaluated using a circular corridor (actimeter) [36].

### Cocaine-induced conditioned place preference (CPP), extinction and reinstatement

The CPP was evaluated in a Plexiglas Y-shaped apparatus (Immetronic) with one arm blocked, located in a soundproof testing room with low luminosity using an unbiased procedure. Two of the three chambers were used for the conditioning and distinguished by different floor textures and wall patterns. The device was connected to an electronic interface for data collection.

Entries and time spent in each chamber were measured, as well as the locomotor activity of the mice during the experiments.

### Operant self-administration of cocaine

The self-administration experiments were conducted in mouse operant chambers (Model ENV307A-CT; Med Associates Inc., Georgia, VT, USA) as previously described [37]. Nose pokes were the operant action that led to delivery of a cocaine infusion.

### Immunocytochemical studies

The procedures for Cre-recombinase (anti-Cre-recombinase antibody; 1:500, mouse, MAB3120, Merck Millipore), and Fos-B (anti-FosB antibody 1:500, rabbit, Ma5-15056, Invitrogen) immunofluorescence experiments and quantification of Fos-B expression are detailed in **supplementary methods**.

### Statistical analysis

All statistical analyses were carried out using GraphPad Prism and Statview softwares. For all experiments, ANOVAs were performed followed, when appropriate, by post hoc test Bonferroni for multiple comparisons, except when data did not pass the normality test and/or when the number of individuals was too low. In those cases, non-parametric tests, such as Mann-Whitney (for 2 groups) or Kruskall-Wallis followed by post-hoc Dunn’s test (for more than two groups) were used. The null hypothesis was rejected at p < 0.05. All the statistical results for principal figures are reported in **Table 1**.

## RESULTS

### Sub-chronic cocaine exposure differentially regulates a subset of plasticity-related miRNAs in the NAc and DS

Several miRNAs have been shown to respond to 1-week cocaine exposure in a broad miRNA sequencing study [31]. Among those, we selected a subset of 14 miRNAs (miR-1, −16, −29b, −31, −32, −124, −125a, −128b, −132, −181a, −212, −221, −223 and Let7) based on their association either with cocaine addiction [25] or with other related brain disorders [24]. We measured the expression levels of these 14 miRNAs in the NAc and DS by RT-qPCR, after a sustained treatment with cocaine mimicking a behavioral sensitization procedure (**Figure 1a**) known to induce modifications in miRNAs expression [31]. The sequences of all miRNA primers used are presented in **Supplementary Figure 1a**. Interestingly, the repeated exposure to cocaine did not significantly alter miRNA levels in the DS but induced an up-regulation of 11 of these 14 miRNAs in the NAc (**Figure 1b** **and** **c**, **left panels**).

**Figure 1:**
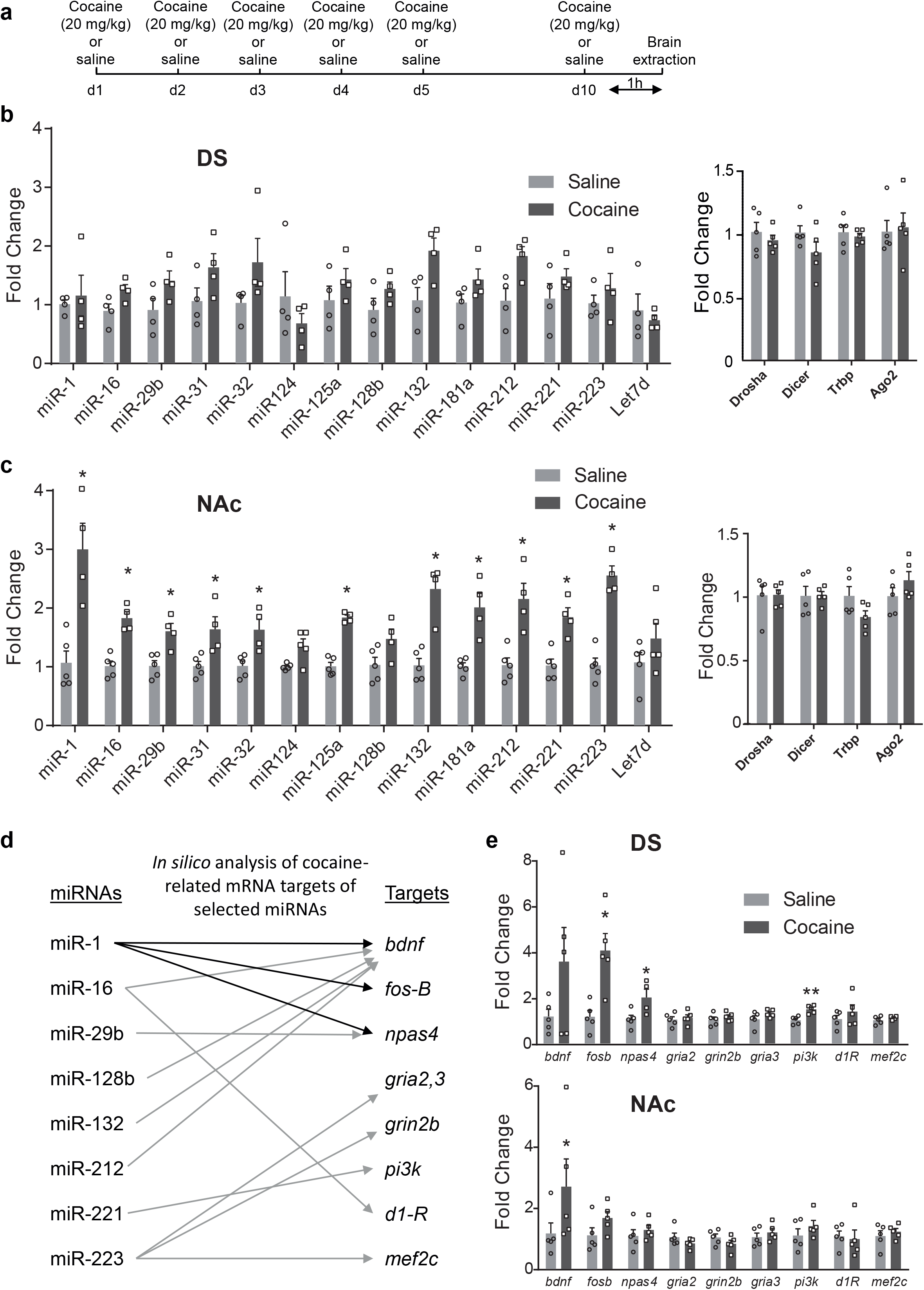
A set of miRNAs is up-regulated in the ventral (NAc), but not the dorsal (DS), striatum after sub-chronic treatment with cocaine. **a)** Schematic representation of the timing of cocaine (20mg/kg) or saline administrations and brain extraction. **b-c)** Expression levels of miRNAs (**left panels**) and mRNAs involved in miRNA processing (**right panels**) were analyzed in the DS (**b**) and the NAc (**c**) one hour after the last injection of cocaine or saline on day 10. **d)** In silico analysis of cocaine-related targets of miRNAs upregulated in the NAc, as indicated in (**c**). **e)** Expression of predicted mRNA targets one hour after the last injection of cocaine or saline on day 10, in the DS (**upper panel**) or the NAc (**lower panel**). **b**) **c**) and **e**) : Mean +/− SEM are represented. Mann-Whitney tests: *p<0.05, **p<0.01 when comparing saline and cocaine groups.

The mRNA expression levels of the key players in miRNA processing, such as *Drosha, Dicer, Tar RNA Binding Protein (TRBP)* and *Ago2*, were not modulated by cocaine (**Figure 1b and c**, **right panels**), indicating that the observed miRNA regulation did not result from alterations in the general miRNA processing machinery at the mRNA level.

In order to predict cocaine-related target genes of the selected miRNAs, we used a combination of databases, including miRBase, Miranda, TargetScan and Tarbase, crossed with literature data on cocaine addiction. We identified 9 potential interesting targets: *bdnf, fos-B, npas4, gria2, grin2b, gria3, pi3K, drd1a* and *mef2c* (**Figure 1d**) and studied expression levels of these mRNAs in the DS and NAc after our protocol of sustained cocaine administration. Among these mRNAs, we only observed a significant increase in the expression levels of *fosB, npas4* and *pi3k* mRNAs in the DS, but not in the NAc (**Figure 1e**).

### Validation of *fosB* and *npas4* mRNAs as targets of miR-1

In most cases, miRNAs interact with the 3′ UTR of target mRNAs to induce mRNA degradation or translational repression [38]. We hypothesized that the absence of detectable up-regulation of *fosB* and *npas4* mRNAs by cocaine in the NAc might be due to cocaine-induced up-regulation of miRNA(s) targeting these mRNAs in this region. Among the selected miRNAs, we focused on miR-1 because the “in silico” prediction indicated that it targeted both *fosB* and *npas4* mRNAs (**Figure 1d**). which are key players in cocaine-induced long-term adaptations [41,42]. We experimentally tested that these potential targets were directly bound by miR-1 (**Figure 2**) using a luciferase reporter assay with constructs containing either *fosB*, *npas*-4 mRNA 3’UTRs (**Figure 2a left panel**), or the *bdnf* 3’UTR as a positive control [39]. We observed a significant decrease of the luciferase activity in HEK cells expressing the pmiRGlo plasmid encoding the 3’UTR of *bdnf, fosB* and *npas4* in the presence of a mimick-miR-1 when compared to the negative miR-control (**Figure 2a**, **right panels**). We thus validated *in vitro* that the 3’UTR of *bdnf*, *fosB* and *npas4* are directly targeted by miR-1.

**Figure 2:**
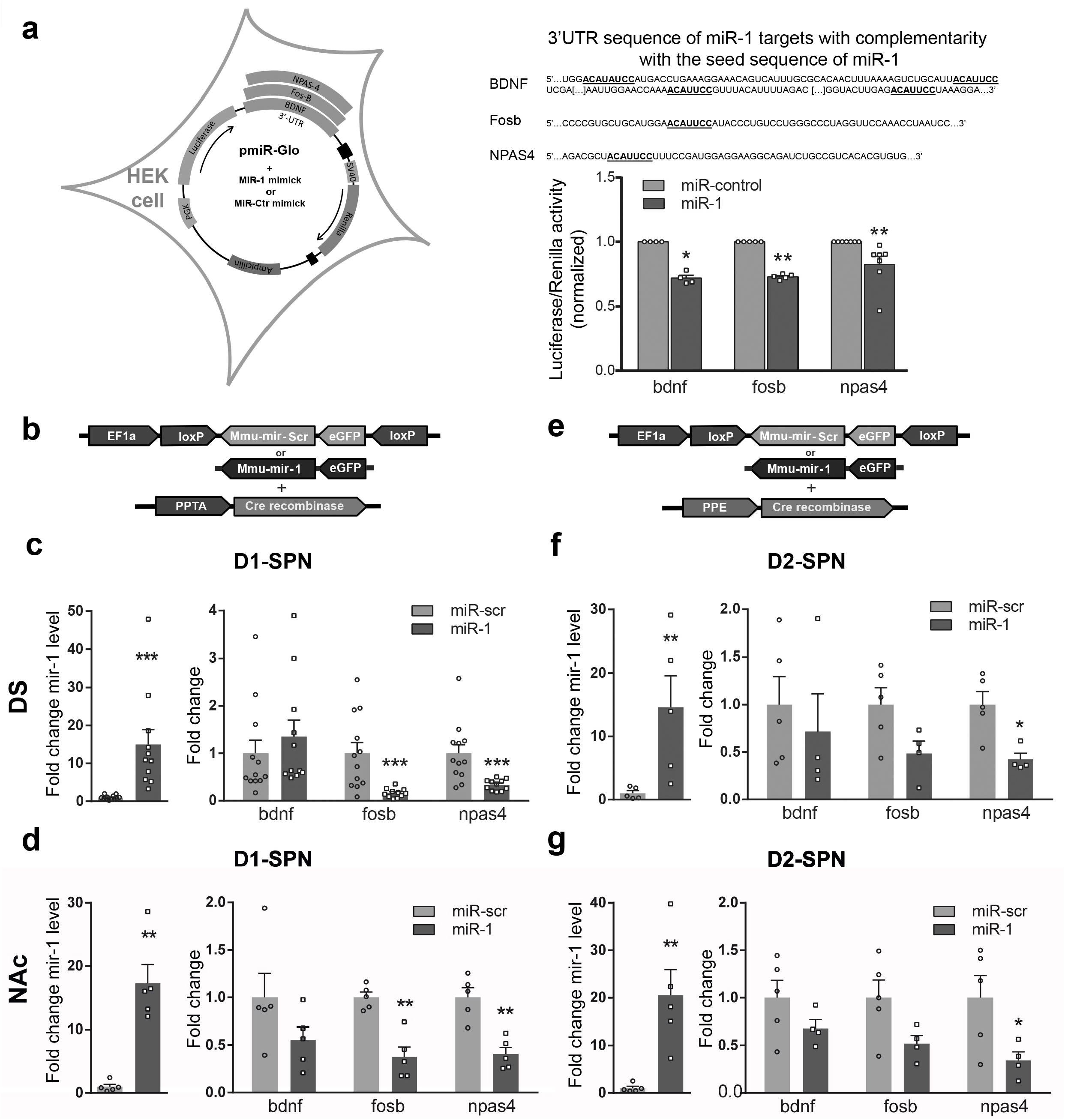
Validation of miR-1 targets. **a) Left panel**: schematic representation of the pmiR-Glo constructs used in HEK cells. **Right Panel**, up: 3’UTR sequences of *bdnf*, *fosB* and *npas4* mRNAs. Bold and underlined nucleotides represent complementarity with the seed sequence of miR-1; lower part: normalized Luciferase/Renilla activity of pmiR-Glo plasmid with *bdnf*, *fosB* and *npas4* 3’UTRs mRNAs in the presence of miRNA mimics for miR-1 (black bars) or miR- control (grey bars). **b-d)** overexpression of miR-1 in D1-SPN. **b)** representation of the combination of AAVs to infect D1-SPNs. **c-d)** normalized fold changes in expression of miR-1 (over miR-scr condition) (**left panels**) and *bdnf*, *fos-B* or *npas4* mRNA (**right panels**) after viral injection of miR-1 or miR-scr sequences in D1-SPNs of the DS (**c**) or the NAc (**d)** of mice. **e-g)** overexpression of miR-1 in D2-SPNs. **e)** representation of the combination of AAVs to infect D2-SPNs. **f-g)** normalized fold changes in expression of miR-1 (over miR-scr condition) (**left panels**) and *bdnf*, *fos-B* or *npas4* mRNA (**right panels**) after viral injection of miR-1 or miR-scr sequences in D2-SPNs of the DS (**f**) or the NAc (**g)** of mice.. **a-g)** Mean +/− SEM are represented. Mann-Whitney tests: *: p<0.05 ; **: p<0.01; ***: p<0.001.

We further aimed to assess whether miR-1 could repress *bdnf*, *fosB* and *npas4* mRNA levels *in vivo.* To this end, we used an AAV-mediated cell-type specific strategy allowing to overexpress miR-1 in either D1R-SPNs or D2R-SPNs. We used a combination of two AAVs, with one expressing miR-1 in a Cre recombinase-dependent manner and the other one expressing Cre under a cell type-specific promoter (*PPTA*::*Cre* and *PPE::Cre*, expressed in D1R-SPNs and D2R-SPNs respectively) as previously validated [32–34] **(see also supplementary Figure 2a-b).** As compared to the control group (injected with an AAV expressing scrambled RNA, Scr) we found a marked increase of miR-1 levels after overexpression of miR-1 in D1R- (**Figure 2b**) or D2R- (**Figure 2c**) in either the DS (**Figures 2b and c**, **upper panels**) or the NAc (**Figures 2b and c**, **lower panels)**, with a concomitant decrease in the expression of *fosB* and *npas4* mRNAs, which was significant in several cases (**Figure 2b-c).** We thus confirmed the ability of miR-1 to repress *fosB* and *npas4* mRNAs expression *in vivo*. Consistent with the preferential expression of *bdnf* mRNA in axons from afferent excitatory cortical neurons [40], and their low levels in SPNs we failed to detect a significant effect of miR-1 overexpression on *bdnf* mRNA levels in any conditions. miR-1 overexpression did not impact on the expression levels of mRNAs involved in miRNA processing, nor levels of miR-206 a miRNA from the same family than miR-1 (**Supplementary Figure 3a and b**).

### Effects of miR-1 overexpression in D1- and D2R-SPNs of the NAc and DS on cocaine-induced locomotor sensitization

Having established the inhibitory role of miR-1 on *fos-b* and *npas4* mRNA expression levels, we investigated the potential role of miR-1 overexpression on behavioral adaptations to cocaine. We decided to perform targeted-overexpression of miR-1 in either D1- or D2-SPNs of the DS and NAc, and first studied the consequences on cocaine-induced locomotor sensitization (15 mg/kg) (**Supplementary Figure 4a-e)**. The overexpression of miR-1 in either D1-SPNs or in D2-SPNs of the NAc or DS did not modify the locomotor sensitization induced by cocaine (**Supplementary Figure 4b-e**).

### Cell-type-specific effects of miR-1 in D1- and D2R-SPNs of the NAc and DS in cocaine-induced conditioned place preference

We then measured the effects of overexpression of miR-1 in cocaine-induced conditioned place preference (CPP), which can be used as a proxy for the rewarding properties of cocaine. Overexpression of miR-1 in D1-SPNs of the DS did not alter cocaine-induced CPP (**Figure 3a**, **left panel).** We then extinguished the CPP by freely exposing the mice to cocaine-paired and unpaired environments after saline injection and observed that miR-1 overexpression in D1-SPNs of the DS did not modify CPP extinction (**Figure 3a**, **middle panel**). In contrast, the reinstatement of cocaine CPP induced by an acute injection of cocaine after the extinction period (15 mg/kg) was prevented by miR-1 overexpression in DS D1-SPNs (**Figure 3a**, **right panel)**.

**Figure 3:**
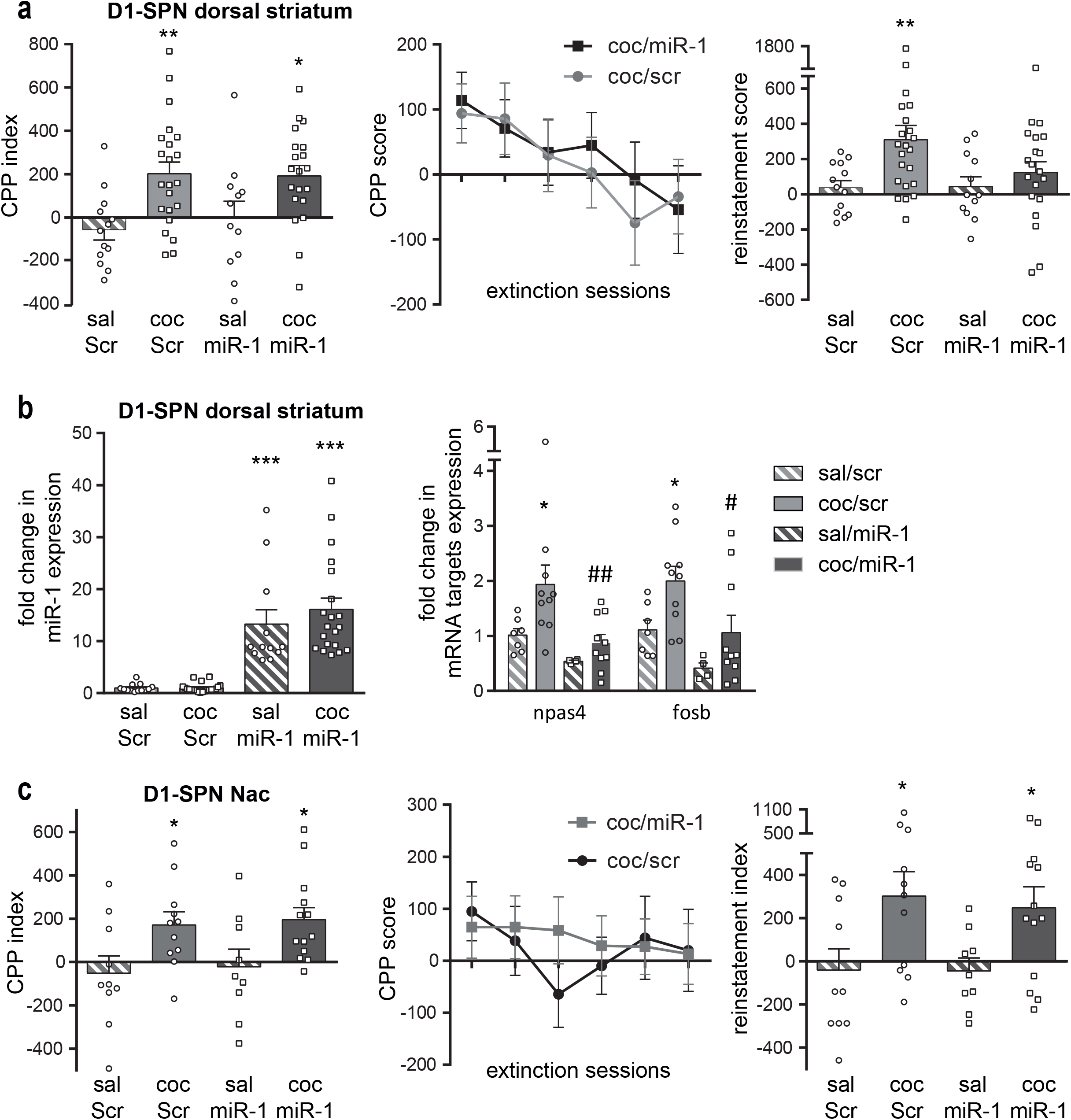
Cocaine-induced conditioned place preference (CPP) and relapse in mice after miR-1 over-expression in D1-SPNs of the DS or NAc. **a-b)** overexpression of miR-1 in D1-SPNs of the dorsal striatum. **a) Left panel:** CPP after miR-1 over-expression in DS D1-SPNs. The CPP index is the difference of CPP score between the pretest and the test. The 2-way ANOVA indicated a significant effect of treatment, no effect of AAV and no interaction. **Middle panel:** extinction sessions for cocaine/miR-Scr and cocaine/miR-1 groups. The 2-way ANOVA indicated a significant effect of session but not of AAV and no interaction. **Right panel:** Cocaine-induced reinstatement of cocaine seeking after extinction in mice with miR-1 over-expression in D1-SPN of the dorsal striatum. The 2-way ANOVA indicated a significant effect of treatment, no effect of AAV and no interaction. Post hoc tests : Bonferroni’s multiple comparison tests: *p<0.05; **p<0.01. **b) Left panel:** expression of miR-1 after cocaine-induced reinstatement of cocaine seeking in mice with miR-1 or miR-Scr over-expression in D1-SPN of the DS. **Right panel:** normalized fold changes of expression levels of *npas4* and *fosB* mRNA after cocaine-induced reinstatement of cocaine seeking in mice with miR-1 over-expression in D1SPN of the DS; The Kruskal-Wallis test on *npas4* and *fosB* mRNAs expression levels indicated a significant group effect. The Post-hoc analysis (Dunn’s multiple comparison test) indicated significant differences between the saline/miR-Scr and the cocaine/miR-Scr groups for both *npas4* and *fosB* mRNAs: *p<0.05 and between the cocaine/miR-Scr and the cocaine/miR-1 groups for both *npas4* and *fosB* mRNAs: #p<0.05; ##p<0.01. **c**) overexpression of miR-1 in D1-SPNs of the NAc. **Left panel :** CPP after miR-1 over-expression in D1-SPN of the NAc of mice. The CPP index is the difference of CPP score between the pretest and the test. **Middle panel:** extinction sessions for cocaine/miR-Scr and cocaine/miR-1 groups. **Right panel:** Cocaine-induced reinstatement of cocaine seeking after extinction in mice with miR-1 over-expression in D1-SPN of the NAc. Bonferroni’s multiple comparison tests: *p<0.05. **a-c)** Mean and SEM are represented.

We measured the levels of miR-1 and its target genes in the DS of these mice 1h after the start of the reinstatement test. As expected we found, high levels of miR-1 in both saline and cocaine miR-1-injected groups, while miR-1 levels were not altered in control groups (**Figure 3b**, **left panel**). miR-1 overexpression in D1-SPNs blunted *fosB* and *npas4* mRNA up-regulation observed in the control scramble group after cocaine-induced reinstatement of CPP (**Figure 3b**, **right panel**). miR-1 overexpression in D1-SPNs of the NAc did not alter either cocaine-induced CPP, its extinction or cocaine-induced reinstatement (**Figure 3c**). As shown in **Figure 4a and b**, the overexpression of miR-1 in D2-SPNs of the DS or NAc had no consequence on cocaine-induced CPP, extinction or cocaine-induced reinstatement. These results indicate that miR1-overexpression in DS D1-SPNs specifically, alters cocaine-induced CPP reinstatement.

**Figure 4:**
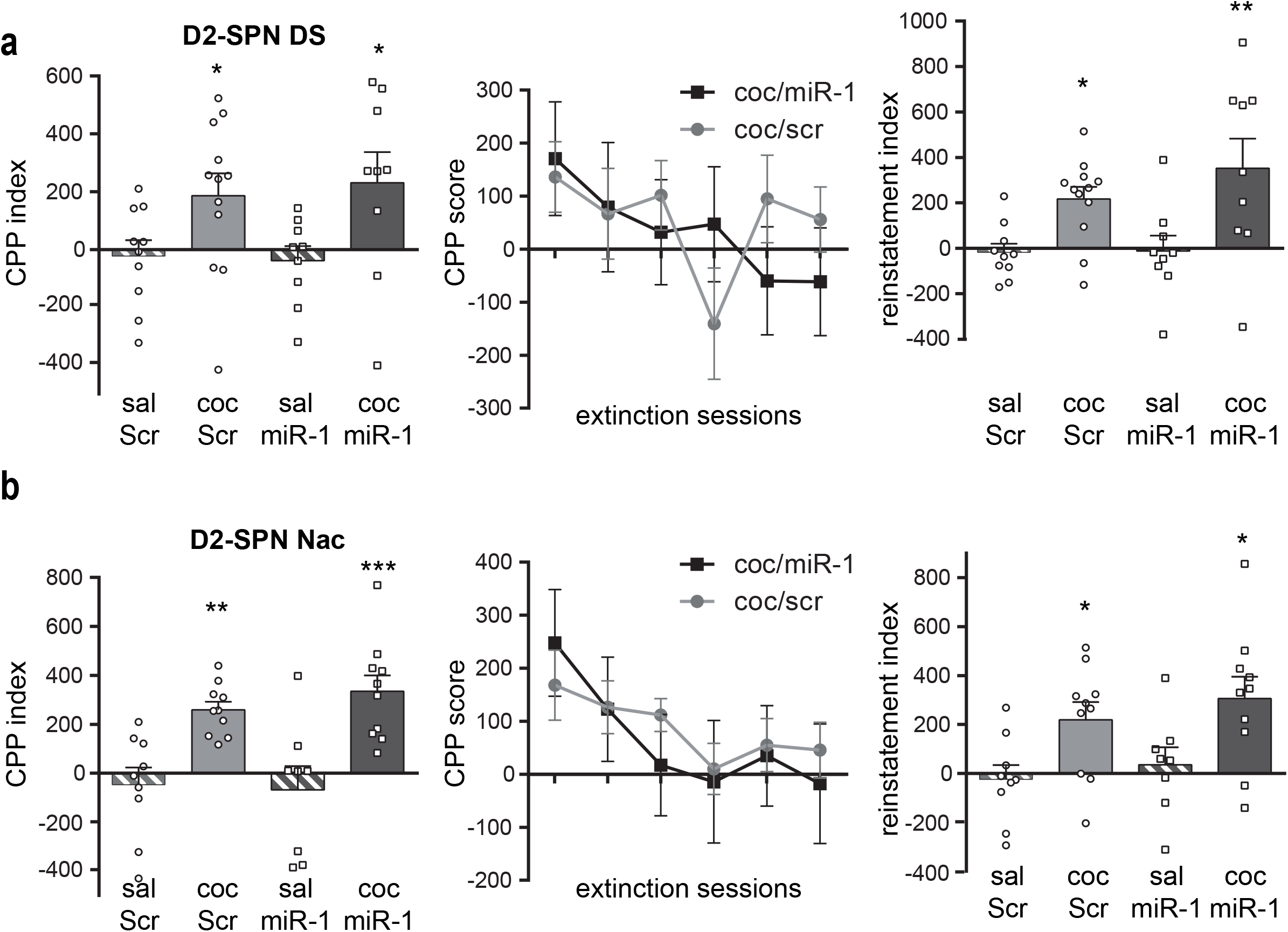
Cocaine-induced Conditioned Place Preference and relapse after miR-1 overexpression in D2-SPN of the dorsal striatum or NAc of mice. a) miR-1 overexpression in D2-SPN of the dorsal striatum (DS) of mice. **Left panel:** CPP. The 2-way ANOVA performed on the CPP index indicated a significant effect of treatment but not of AAV and no interaction. **Middle panel:** extinction sessions for coc/ Scr and coc/miR-1 groups. The 2way ANOVA indicated a significant effect of session but not of AAV and no interaction. **Right panel:** Cocaine-induced reinstatement of cocaine seeking after extinction. The two-way ANOVA performed on the reinstatement index (difference of CPP score between the extinction and the reinstatement test) indicated a significant effect of treatment but not of AAV and no interaction. b) miR-1 overexpression in D2-SPN of the NAc of mice. **Left panel:** CPP. The two-way ANOVA indicated a significant effect of treatment but not of AAV and no interaction. **Middle panel:** extinction sessions for coc/Scr and coc/miR-1 groups. The 2-way ANOVA indicated a significant effect of session but not of AAV and no interaction. **Right panel:** Cocaine-induced reinstatement of cocaine seeking after extinction. The two-way ANOVA indicated a significant effect of treatment but not of AAV and no interaction. **a-b)** Mean +/− SEM are represented. Post-hoc Bonferroni’s multiple comparison tests *p<0.05; **p<0.01; ***p<0.001.

### Cocaine self-administration, extinction and reinstatement after miR-1 overexpression in D1- or D2-SPNs of the DS of mice

We then investigated whether miR-1 overexpression in D1-SPNs of the DS altered cocaine self-administration. MiR-1 overexpression in DS D1-SPNs did not modify the acquisition of self-administration, the motivation for cocaine under a progressive ratio of reinforcement (**Figure 5a-b**), nor the extinction of this behavior (**Figure 5c**). Furthermore, the number of days required for acquisition of cocaine self-administration, or for achievement of extinction was similar between groups. However, miR-1 overexpression in D1-SPNs of the DS induced a significant increase in cue- induced reinstatement of cocaine seeking (**Figure 5d**), a result that contrasted with the blockade of cocaine-induced reinstatement of CPP (see **Figure 3a**). Interestingly, when analyzing FosB expression after cue-induced reinstatement of cocaine seeking, we found a slight decrease in the DS and an increase in the NAc core after miR-1 overexpression in the D1-SPNs of the DS (**Figure 5e**), a result that suggests the existence of crosstalk between the DS and NAc.

**Figure 5:**
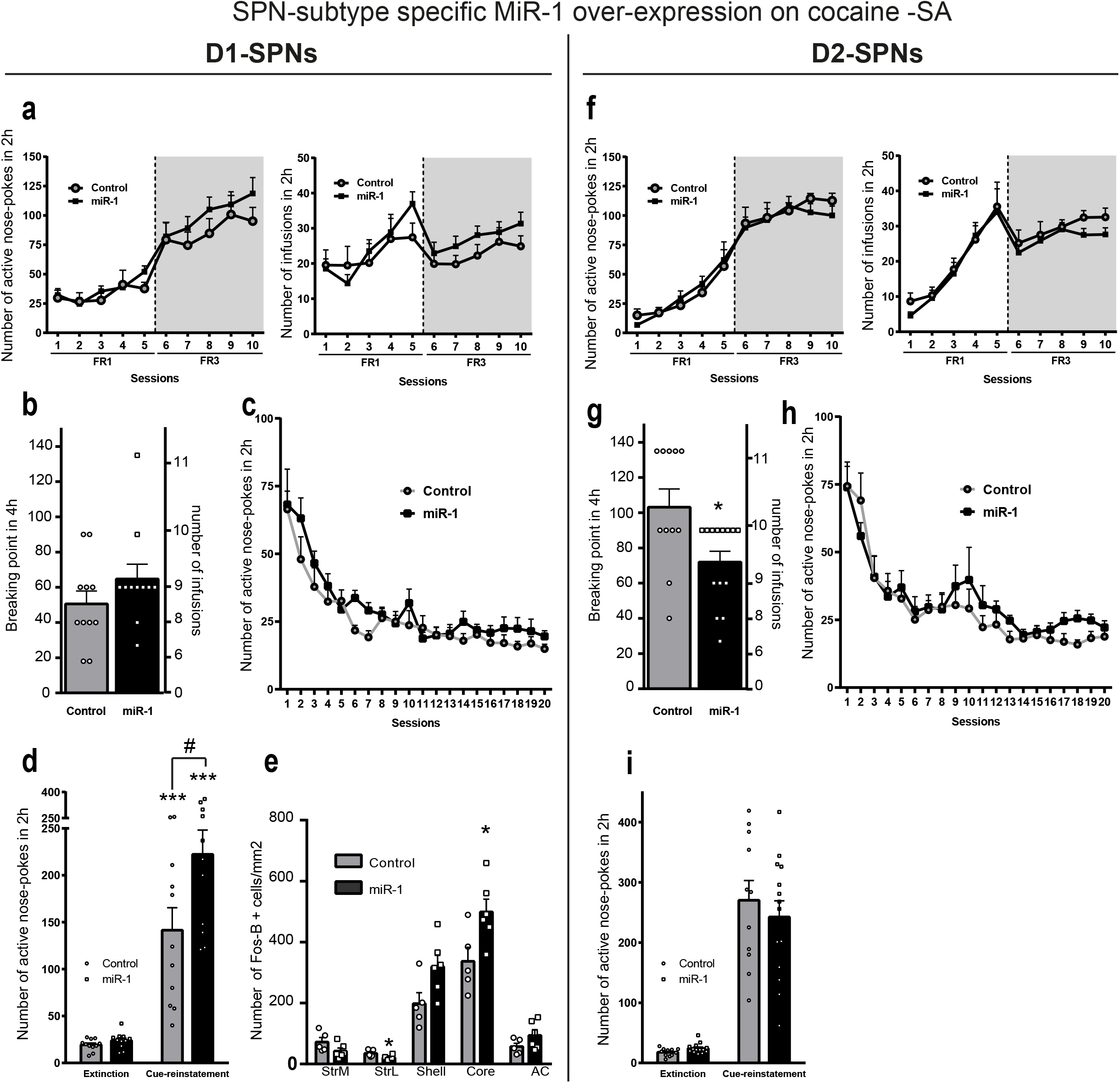
Cocaine Self-Administration and relapse after extinction in mice with miR-1 overexpression in D1- or D2-SPNs of the dorsal striatum. **a-e)** miR-1 overexpression in D1- SPNs of the dorsal striatum. **a**) Cocaine Self-Administration. **Left panel**: active nose-pokes during the acquisition of cocaine (0.5 mg/kg/infusion)-induced self-administration in mice with (n=11) or without (n=11, scr) miR-1 overexpression in the D1-SPNs of the DS. The 2-way ANOVA indicated no effect of miR-1 overexpression, a significant effect of session, and no interaction. **Right panel:** number of cocaine infusions received during the acquisition of cocaine -induced self-administration. The 2-way ANOVA indicated no effect of miR-1 overexpression), a significant effect of session and no interaction. **b**) Breaking point for cocaine (0.5mg/kg/infusion)-induced self-administration under a progressive ratio . The Mann-Whitney test indicated no AAV effect. **c**) Active nose-pokes during the extinction sessions. The 2-way ANOVA indicated no effect of miR-1 overexpression, a significant effect of session and no interaction. **d**) Cue-induced reinstatement of cocaine seeking after extinction (miR-1 overexpression in the D1-SPNs of the DS). The 2-way ANOVA indicated a significant effect of miR-1 overexpression, a significant effect of cue-reinstatement and a significant interaction. Post-hoc Bonferroni’s multiple comparison test ***: p<0.01 (extinction vs cue-reinstatement); #: p<0.05 (scr vs miR-1). **e**) Quantification of expression levels of the Fos-B protein 24h after cue-induced reinstatement of cocaine seeking. The MannWhitney tests indicated significant differences between the scr and the miR-1 groups in the lateral part of the DS (Strl) and the Core of the NAc: *p<0.05. **f-j)** miR-1 overexpression in D2- SPNs of the dorsal striatum. **f**) Cocaine Self-Administration. **f**) Cocaine Self-Administration. **Left panel**: active nose-pokes during the acquisition of cocaine (0.5 mg/kg/infusion)-induced self-administration in mice with (n=14) or without (n=11, scr) miR-1 overexpression in the D2-SPNs of the DS. The 2-way ANOVA indicated no effect of miR-1 overexpression, a significant effect of session, and no interaction). **Right panel:** number of cocaine infusions received during the acquisition of cocaine -induced self-administration. The 2-way ANOVA indicated no effect of miR-1 overexpression, a significant effect of session and no interaction. **g**) Breaking point for cocaine (0.5mg/kg/infusion)-induced self-administration under a progressive ratio (miR-1 overexpression in the D2-SPNs of the DS). The Mann-Whitney test indicated a significant effect of miR-1 overexpression (p=0.018). **h**) Active nose-pokes during the extinction sessions after miR-1 overexpression in the D2-SPNs of the DS. The 2-way ANOVA indicated no effect of miR-1 overexpression, a significant effect of session and no interaction. **i**) Cue-induced reinstatement of cocaine seeking after extinction. The 2-way ANOVA indicated no effect of miR-1 overexpression, a significant effect of cue-reinstatement and no interaction. Post-hoc Bonferroni’s multiple comparison test ***: p<0.01 (extinction vs cue-reinstatement). **j**) Quantification of expression levels of the Fos-B protein 24h after cue-induced reinstatement of cocaine seeking. **a-i)** Mean +/− SEM are represented. **e**) and **j**) abbreviations : StrM: striatum dorsal median; StrL: striatum dorsal lateral; Shell: NAc shell; Core: NAc core; AC: anterior cingulate cortex.

MiR-1 overexpression in D2-SPNs of the DS did not alter the acquisition of cocaine-SA (**Figure 5f)** nor its extinction (**Figure 5h)** or cue-induced reinstatement of cocaine seeking (**Figure 5i)**. Interestingly, it significantly reduced the breaking point for cocaine –SA under a progressive ration schedule of reinforcement (**Figure 5g**), suggesting a reduced motivation for cocaine-SA.

## DISCUSSION

Our data show that miR-1 is up-regulated in the NAc, but not the DS, by a 10-day regimen of cocaine and that at least three mRNAs encoding proteins associated with cocaine-induced neuronal plasticity, *fosB, npas4* and *bdnf* are targeted by miR-1. We also demonstrate that overexpression of miR-1 in D1-SPNs of the DS decreases cocaine-induced reinstatement of cocaine seeking in a CPP paradigm, but increases cue-induced reinstatement of cocaine seeking in a SA protocol. In addition, overexpression of miR-1 in D2-SPNs of the DS reduces the breaking point for cocaine-SA under a PR schedule of reinforcement.

Multiple miRNAs were reported to be modulated by cocaine [26]. In a miRNA profiling study, a panel of miRNAs was either up- or down-regulated by prolonged cocaine exposure [43]. Induction of miR-132 and miR-212 occurred in the DS after extended access to cocaine self-administration, and overexpression of miR-212 reduced compulsive cocaine intake in rats through CREB, MeCP2 and BDNF signaling [44,45]*. In silico* genome-wide sequencing identified miR-495 among miRNAs whose targets are enriched in an addiction-related gene (ARG) database. Lentiviral-mediated overexpression of miR-495 in the NAc suppressed the motivation to self-administer and seek cocaine [46].

Deficiency of Ago2 in D2-SPNs greatly influenced a specific group of miRNAs, and reduced approximately by half the expression of miR-1 [27], suggesting that miR-1 is expressed in both types of SPNs in this structure. Similarly to our study, Hollander et al. (2010) showed that overexpression of miR-1 in the DS, failed to alter cocaine intake, but cell-type-specificity and reinstatement of drug seeking were not investigated [44]. Herein, we provide evidence that miR-1 targets the mRNA encoding the transcription factor FosB, the spliced version of which, ΔFosB, persistently accumulates in D1-SPN of the NAc after chronic psychostimulant exposure, and plays a key role in cocaine addiction [47]. This D1-SPN cell-type specific ΔFosB expression induced by psychostimulant was further confirmed in reporter lines [48,49] or using ribosomal tagging approaches [50]. Furthermore, ΔFosB overexpression in D1-SPN increases locomotor activity and rewarding effects of low doses of cocaine [51], as well as cocaine self-administration [52]. Since miR-1 reduces the expression *fosb* mRNAs, which encodes both FosB and ΔFosB, it was surprising to observe that miR-1 overexpression in D1-SPNs of the NAc failed to alter cocaine-induced locomotor sensitization and CPP. However, the involvement of ΔFosB in cocaine’s effect seems highly dependent on the environment. For example, chronic cocaine treatment elevates ΔFosB protein levels in the NAc of isolated rats, but not in enriched condition due to an already elevated basal accumulation of ΔFosB [53]. Furthermore, the facilitating effects of ΔFosB overexpression in D1-SPN on cocaine-induced behaviors were seen in individually housed animals [52], In addition, overexpression of ΔFosB in the NAc shell of pair-housed rats increases operant responding for sucrose when motivated by hunger, but decreases responding in satiated animals, in cocaine self-administration and cocaine-induced relapse. In our study, the mice were housed five per cage, and we did not observe a significant increase of *fosB* mRNAs in the NAc upon sustained cocaine administration, similarly to previous studies in rats [53]. The absence of effect of miR-1 overexpression in D1-SPNs of the NAc on cocaine-induced locomotor sensitization and CPP might thus be due to our environmental conditions. Alternatively, we cannot exclude that ΔFosB overexpression previously observed in D1-SPNs of the NAc [47–49] results from an accumulation of the stable protein rather than from de novo gene expression, or that a partial decrease induced by miR-1 overexpression may not be sufficient to prevent the behavior.

Another validated target of miR-1 was *bdnf* mRNA encoding brain-derived neurotrophic factor (BDNF), which also plays a key role in cocaine-induced behaviors [54]. In particular, data from several labs suggest a facilitating effect of BDNF protein in the NAc on cocaine-induced CPP and cocaine-primed reinstatement [55,56]. However, the main source of BDNF within the striatum, including the NAc, arises from prefrontal cortical neurons release [57], which mediates the majority of BDNF effects on cocaine-related behaviors [58]. By contrast, *bdnf* mRNA levels are very low in the striatum under basal conditions, as observed in our RT-qPCR data. Altogether, these observations can explain the lack of effect of miR-1 overexpression in *bdnf* mRNA levels *in vivo*, despite a clear demonstration that miR-1 targets its 3’UTR region *in vitro*. Further experiments, which are out of the scope of our study, could include overexpression of miR-1 in prefrontal cortical neurons.

Finally, we demonstrate that *npas4*, is a target of miR-1 both *in vitro* and *in vivo*. *npas4* encodes an activity-dependent transcription factor that plays a key role in neuronal plasticity and is induced in the NAc by cocaine [42]. NPAS4 expression is higher in D1- than in D2-SPN in the NAc after cocaine-induced CPP [59] and NPAS4 positive cells co-express cFos [42], another activity-related immediate early gene, which is selectively induced by cocaine in D1-SPN [60]. Conditional deletion of *Npas4* in the NAc significantly reduced cocaine-induced CPP and delayed the acquisition of cocaine self-administration, without affecting cue-induced reinstatement of cocaine seeking [42]. In our study, miR-1 overexpression reduced but did not abolish *npas4* mRNA, a result that could explain its lack of effect in D1-SPN of the NAc on cocaine-induced CPP.

From a behavioral point of view, we found a specific, albeit apparently opposite, effect of miR-1 overexpression in D1-SPN of the DS on cocaine-induced reinstatement of cocaine CPP and cue- induced reinstatement of cocaine seeking in a self-administration paradigm. Thus, while miR-1 overexpression in DS D1-SPN prevented cocaine-induced reinstatement of CPP, it potentiated cue-induced reinstatement of self-administration. These differential effects on cocaine-induced reinstatement of a cocaine-induced CPP and on cue-induced reinstatement of cocaine-induced SA could be explained by differences in cocaine doses and routes of administration used in the two protocols, the time period of extinction and by the fact that the reinstatement of cocaine seeking is induced by a priming dose of cocaine for CPP and by a cue previously associated with cocaine (thus in cocaine-free state) in SA. This differential effect may be due to differences in neuronal circuits involved in each behavior (see [61] and [62] for reviews). [63]. It would have been interesting to evaluate the effect of miR-1 overexpression in D1-SPN on cocaine-induced reinstatement after SA and extinction, but cocaine priming does not induce reliable reinstatement of cocaine seeking in strain of mice (C57Bl6) [64] used in this study, contrasting with Swiss albino mice [65], In addition, in SA, cue-induced reinstatement is mediated by dopaminergic projections from the VTA to the basolateral amygdala (BLA) and glutamatergic projections from the prefrontal cortex to the BLA and NAc, while cocaine-induced reinstatement is independent from the BLA [61]. Thus, the differential effects observed after miR-1 overexpression in D1- SPN of the DS in the operant versus non-operant behaviors, could be due to these specific circuits onto BLA.

Finally, we found a decrease in breaking point for cocaine-SA under a PR schedule of reinforcement induced my miR-1 overexpression in D2-SPNs of the DS, suggesting a reduction of motivation for cocaine-SA. There are very few examples in the literature for specific reduction of motivation for cocaine-SA. D2-SPNs of the DS are inhibitory neurons projecting to the external part of the Globus Pallidus (GPe), which itself contains inhibitory neurons projecting to the subthalamic nucleus (STN). The inhibition of D2-SPN activity thus results in an inhibition of STN activity. Of interest, either lesion or inhibition of STN neurons can lead to a reduction of the breaking-point for cocaine-SA without altering SA under a fixed ratio schedule (66) much like what we observed in the present study. It is thus possible that miR-1 overexpression in D2-SPNs of the DS exerts its reducing effect on motivation for cocaine via an action of the GPe-STN pathway.

In conclusion, our data show a specific and distinct role of miR-1 in cocaine- and cue-induced relapses of two different cocaine-dependent behaviors after an extinction period and on motivation for cocaine-SA. This contributes to the dissection of the mechanisms underlying addictive behaviors. Our results show the cell type-specific and region-specific role of miR-1 and underlines differences between cocaine- and cue-induced reinstatement in non-operant and operant paradigms, and thus highlights the complexity of the control of addictive behaviors and emphasizes the importance of regional cell-type specific investigations.

## FUNDING

This work was supported by Fondation pour la Recherche Médicale (*DEQ20150734352 to JC*), Centre National pour la Recherche Scientifique (CNRS, B.F., A.G., V.K., P.P., P.V., J.C.), Institut National pour la Santé et la Recherche Médicale (INSERM; B.F., A.G., V.K., P.P., P.V., J.-A.G, J.C), Sorbonne Université, Faculté des Sciences et Ingénierie (B.F., A.G., V.K., P.P., P.V., J.-A.G, J.C) Labex Biopsy *Investissements d’Avenir*, ANR-11-IDEX-0004-02 (B.F., A.G., V.K., P.P., P.V., J.-A.G, J.C), the Spanish Ministerio de Economía y Competitividad-MINECO (#SAF2017-84060-R-AEI/FEDER-UE; E.M.-G., L.D.-R., R.M.), the Spanish Instituto de Salud Carlos III, RETICS-RTA (#RD12/0028/0023; E.M.-G., L.D.-R., R.M.), the Generalitat de Catalunya, AGAUR (#2017-SGR-669; E.M.-G., L.D.-R., R.M.), ICREA-Acadèmia (#2015) and the Spanish Ministerio de Sanidad, Servicios Sociales e Igualdad, Plan NAcional Sobre Drogas (#PNSD-2017I068) to R.M., Fundació La Marató-TV3 (#2016/20-30) to E.M.-G, *Fondation pour la Recherche Médicale* (FRM#DPA20140629798) and ANR (ANR-16-CE16-0018) to J.A.G.

## Financial Disclosures

The authors declare no competing interests.

## AKNOWLEDGEMENTS

The authors thank the participation of Marie-Charlotte Allichon for some viral injections, Mélanie Marias and Guillaume Ubiema for immunohistochemcal studies in vivo. We would particularly like to thank the whole team of Jocelyne Caboche for its encouraging support and help to the work.

## Author contributions

B.F. an J.C. designed the study and wrote the manuscript. BF performed the viral injections, cocaine-induced locomotor sensitization and conditioned place preference experiments, RT-qPCR experiments and *in vitro* experiments. A.G. performed RT-qPCR experiments and participated in the writing of the manuscript. E.M.G E.S.M., and L.D.R. performed the cocaine-self-administration experiment. V.K. M.M. and M.C.A participated in the RT-qPCR and immunofluorescence experiments. P.P. performed most of the in vitro experiments and participated in the RT-qPCR experiments. A.A performed viral infections and immunofluorescence experiments for tracing D1 and D2 striatal pathways. R.M. designed the cocaine self-administration experiments. P.V., J.A.G. and R.M. participated in the writing of the manuscript.

## Notes

### Competing Interest Statement

The authors have declared no competing interest.

