## Supplemental materials for "Cell type- and region-specific modulation of cocaine seeking by micro-RNA-1 in striatal projection neurons"

***Supplementary Figure 1, Forget et al.***

**a**

| **miRNAs** | **RT-primers** | **Forward primers** | **Reverse primers** |
| --- | --- | --- | --- |
| SnoU6 | CACGGAAGCCCTCACACCGTGTCGTTC | GCTTCGGCAGCACATATACTAAAT | CTCACACCGTGTCGTTCCA |
| mir-1 | CTCGACAAACGTCTAAATGCTTACATACTTC | CCAGTGGAGCTGGAATGTAAAGAA | GCTCGACAAACGTCTAAATGCT |
| mir-16 | CCTTTGAGGTTGGTACTACGGCGCCAATAT | GTGCAGTAGCAGCACGTAAAT | ACCTTTGAGGTTGGTACTACGG |
| mir-29b | CAAGCCCTAGTATTCCTCGACAACACTGATT | AGCTCCGTATAGCACCATTTGAAA | GCAAGCCCTAGTATTCCTCGAC |
| mir-31 | ACGAACGGCGTCATGTAACAGCTATGC | ACTCCGGCAAGATGCTGG | GAGTACGAACGGCGTCATGTAA |
| mir-32 | TCTACCAAACGCCAAACCTGCAACTTAG | ACGGTGGAGTATTGCACATTACTA | TGTTCTACCAAACGCCAAACCT |
| mir-124 | CACCGTTCCGCGCCGTCGGTGGGCCATT | CATACCTAAGGCACGCGG | GTCGGTGGGCATTCACC |
| mir-125a | CCTCACAACGATTCCACAAGCACAGGTTA | GTTGATTCTCCCTGAGACCCTTTA | GTCCTCACAACGATTCCACAAG |
| mir-128b | CATCCGGCTATGCGTATTGGAAAGAGACC | TGGCGTCACAGTGAACCG | CCTCATCCGGCTATGCGTATTG |
| mir-132 | CGGTACTGGATGACCAGTAGCCTAACGACCAT | TGGTCGATAACAGTCTACAGCC | GGTACTCGATGACCAGTAGCCT |
| mir-181a | TGAACGTAACTCGAATCCGTACTCACCG | CGTCCAACATTCAACGCTGTC | CGTGAACGTAACTCGAATCCGT |
| mir-212 | GTCGGAGGAACGTAGAAGTGGCCGTG | CTCACGTAACAGTCTCCAGTCA | ATCGTCGGAGGAACGTAGAAGT |
| mir-221 | CCAGTCCACGATAGTCCCTTGAAACCCAG | AGTAGGAAAGCTACATTGTCTGCT | AACCAGTCCACGATAGTCCCTT |
| mir-223 | AGGCGTCGAGCTTAATGTCGGGGTATTTG | CGTAGCCCTGTCAGTTTGTCAA | TCTAGGCGTCGAGCTTAATGTC |
| let-7d | GCTCAGACAGAAGTCACACTGAGCACTATG | GCGGGCGGAGAGGTAGT | GTCACACTGAGCACTATGCAAC |
| mir-206 | ACGAGTTTAGAGCCGGATAGCCACACAC | GCGTTGTCTGGAATGTAAGGAAGT | TGACGAGTTTAGAGCCGGATAG |

**b**

| **Gene** | **Forward primers** | **Reverse primers** |
| --- | --- | --- |
| Rpl38 | AGGATGCCAAGTCTGTCAAGA | TCCTTGTCTGTGATAACCAGGG |
| Ppia | CAAATGCTGGACCAAACACAAACG | GTTCATGCCTTCTTTCACCTTCCC |
| HPRT | CCTCACTGCTTTCCGGAGCGG | GGACTGCGGGTCGGCATGA |
| TRBP | AGTCTGAGTGCAACCCCG | CTCTTGGGTCACCATGTACTCT |
| Dicer | CCATGGCAACAAGAAGCAATTC | GCCAGTGTTCAAGCACACAAT |
| Ago2 | AACCTGAGAAATGCCCTCGG | ATCAAACACTGGCTTCCGGT |
| Drosha | GCCTGTGTGGATTCGCTGTA | AATCACGGAGCCTGCTTGTC |
| BDNF | CAGGTGAGAAGAGTGATGACC | ATTCACGCTCTCCAGAGTCCC |
| FosB | GTGAGAGATTTGCCAGGGTC | AGAGAGAAGCCGTCAGGTTG |
| NPAS4 | AGGGTTTGCTGATGAGTTGC | CCCCTCCACTTCCATCTTC |
| Gria2 | GGGAAGTAAGGAAAAGACCAGTGC | CATTGCCAAACCAAGGCCCC |
| Grin2b | CCTGTGTGAGAGGAAATCTCGG | GCTGGATGCCGGGGATAGAA |
| Gria3 | CCCATGCTCTTGTCAGCTTCGT | AGTCCACCTATGCTGATGGTGT |
| Pi3K | ACAGATCGACTTCAGGCGGAAAC | GGGCCACCAGGACAAACTCC |
| DRD1 | AAACCCACAAGCCCCTCTGA | GATGAATTAGCCCACCCAAAC |
| Mef2c | TGCTGGTCTCACCTGGTAAC | ATCCTTTGATTCACTGATGGCAT |

**Supplementary figure 1: sequences of the primers used for RT-QPCR analysis**

**a**: sequences of the RT, Forward and Reverse primers used to study the expression of specific miRNAs by RT-QPCR.

**b**: sequences of the Forward and Reverse primers used to study the expression of specific mRNAs by RT-QPCR.

***Supp Figure 2, Forget et al.***

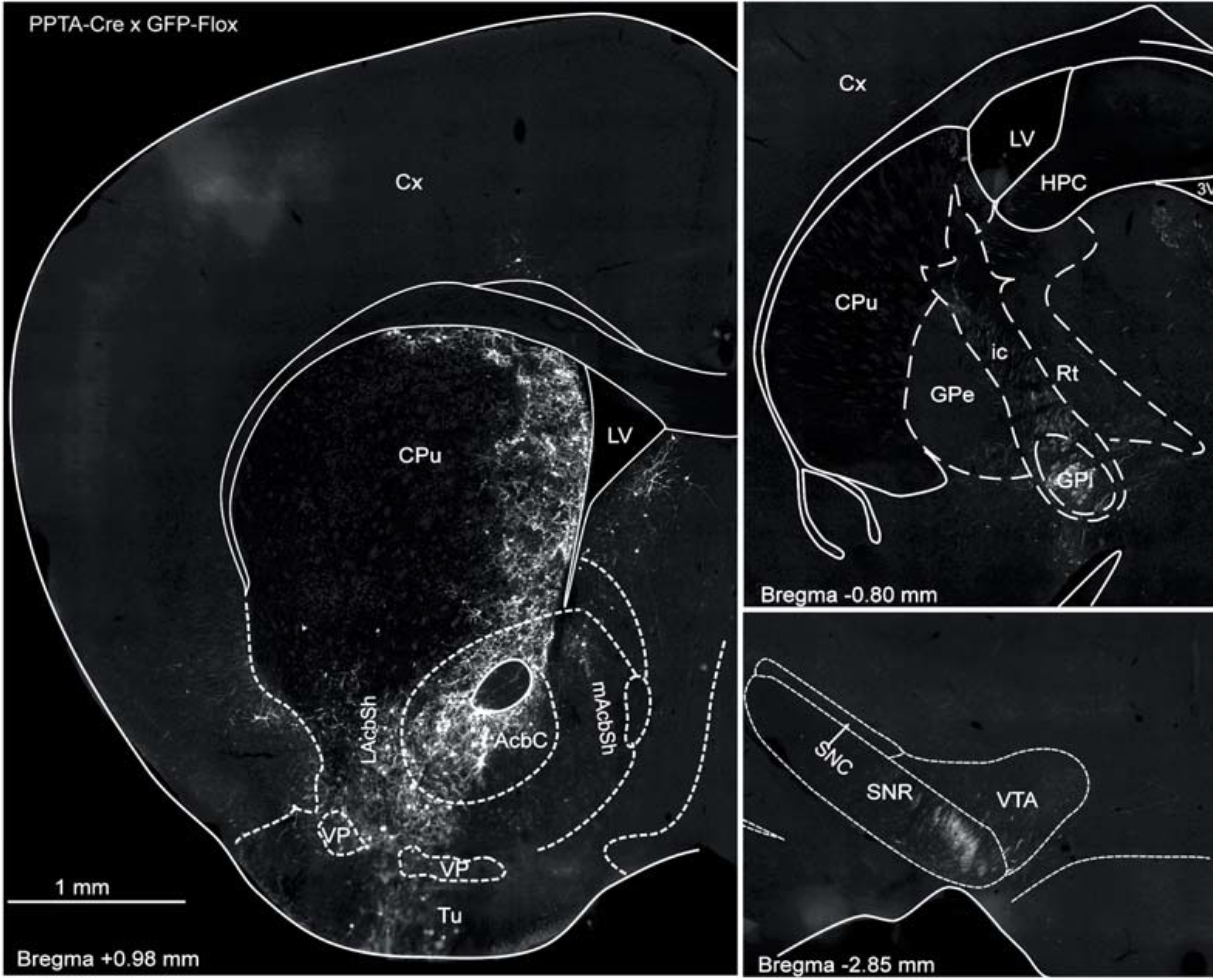

PPTA-Cre x GFP-Flox

a

a

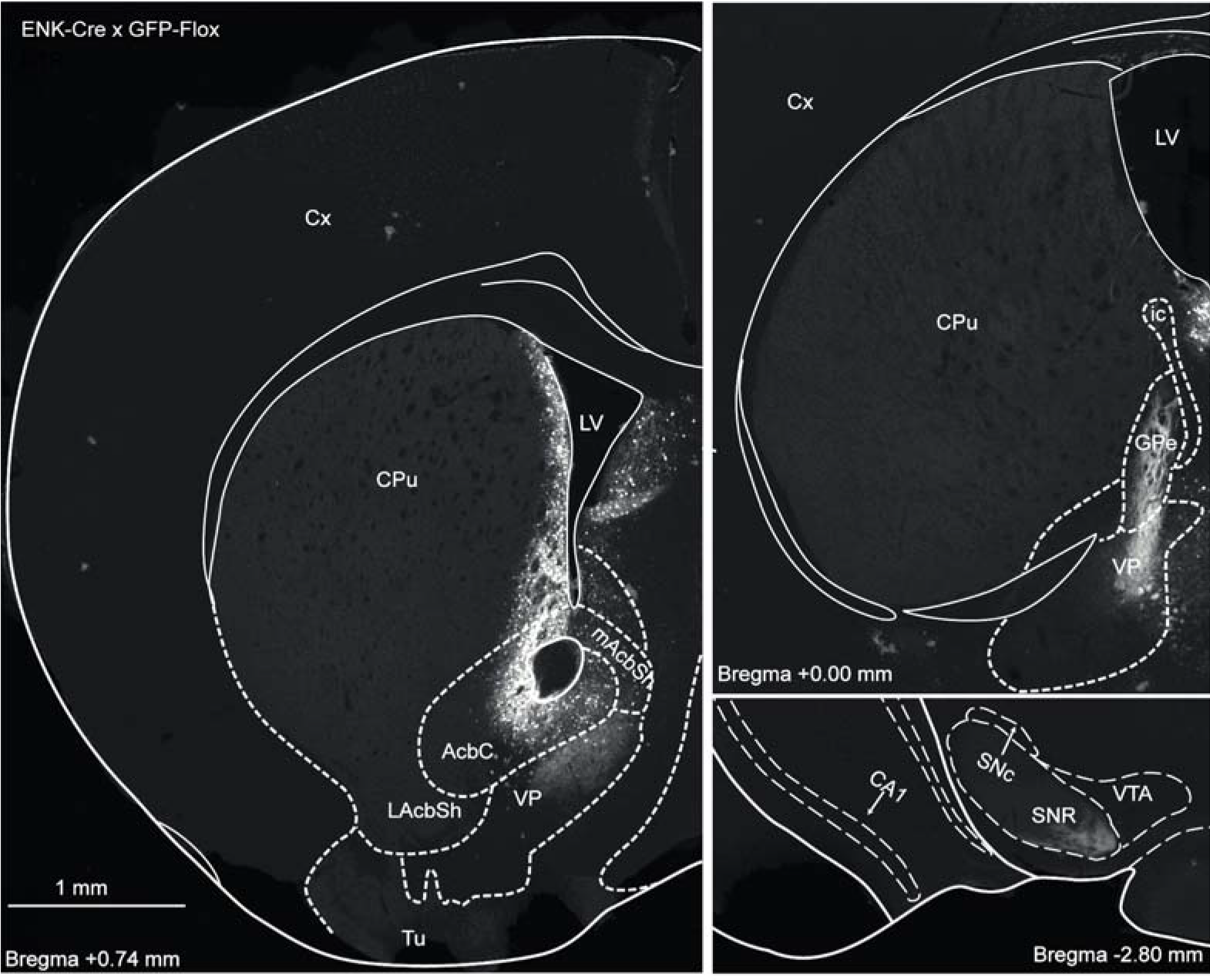
b

PPE-Cre x GFP-Flox

**Supplementary figure 2: validation of the viral strategy for expression in D1- or D2-SPNs.** C57Bl/6J mice were bilaterally injected with a combination of **a:** AAVs (AAV9-PPTA-Cre and AAV9-GFP^flox^) that allows GFP expression in DRD1-expressing neurons. **left panel :**GFP fluorescence from AAV-infected infected neurons in the striatum. **right panel :** depicts GFP fluorescence in projection structures from the direct pathway (DRD1-expressing neurons), such as the GPi, VP, and SNR. **b)** AAVs (AAV9-PPE-Cre and AAV9-GFP^flox^) that allows GFP expression in DRD2-expressing neurons. **left panel :** GFP fluorescence from AAV-infected infected neurons in the striatum. **right panel :** depicts GFP fluorescence in projection structures from the indirect pathway (DRD2-expressing neurons), such as the GPe, VP, and SNR. Abbreviations: LV lateral ventricle; Cx cortex; CPu *caudate putamen* (dorsal striatum); AcbC core of the *nucleus accumbens*; mAcbSh medial shell of the *nucleus accumbens*; LAcbSh lateral shell of the *nucleus accumbens*; VP ventral *pallidum*; Tu olfactory tubercles; GPe external *globus pallidus*; GPi internal *globus pallidus*; ic internal capsule; RT mesencephalic reticular projection; SNR *substantia nigra pars reticulata*; SNC *substantia nigra pars compacta*; VTA ventral tegmental area. CA1 layer of the hippocampus.

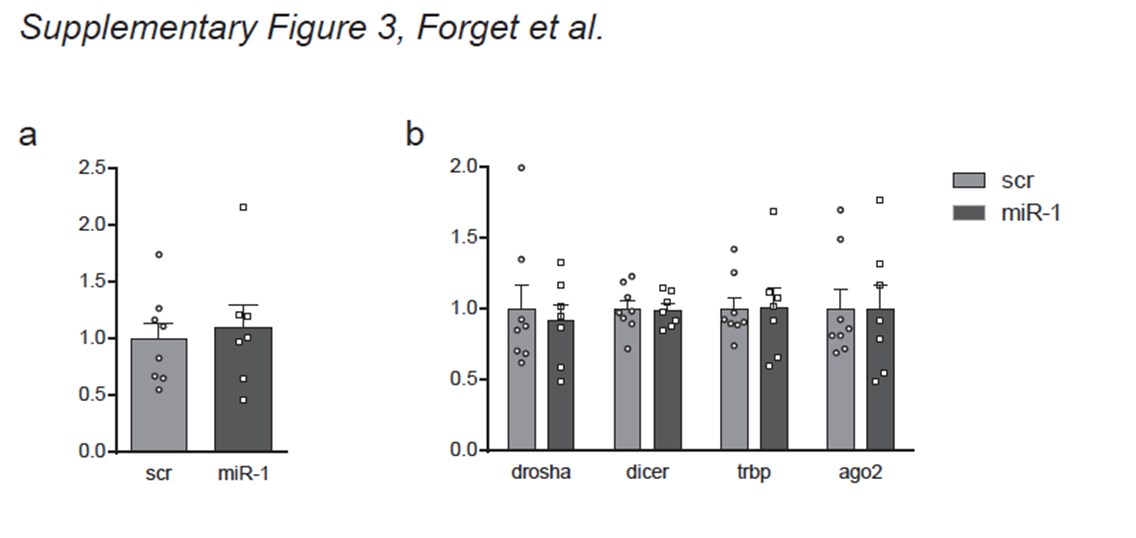
**Supplementary Figure 3: miR-1 over-expression in the Dorsal Striatum did not modify the levels of miR-206 (miR-1 family) or of mRNAs involved in miRNA processing.**

**a**: fold change in expression of miR-206 in the dorsal striatum after miR-1 or miR-Scr overexpressions in the same structure. The Mann-Whitney test revealed no significant difference between the miR-1 and scr groups (U=26.5, p=0.89, NS). Mean and SEM are represented.

**b**: fold change in expression of mRNAs involved in miRNA processing after miR-1 or miRScr over-expressions in the dorsal striatum. The Mann-Whitney tests revealed no significant effect of miR-1 overexpression (drosha: U=28, p>0.9; dicer: U=24.5, p=0.72; trbp: U=26, p=0.84; ago2: U=27, p=0.93). Mean and SEM are represented.

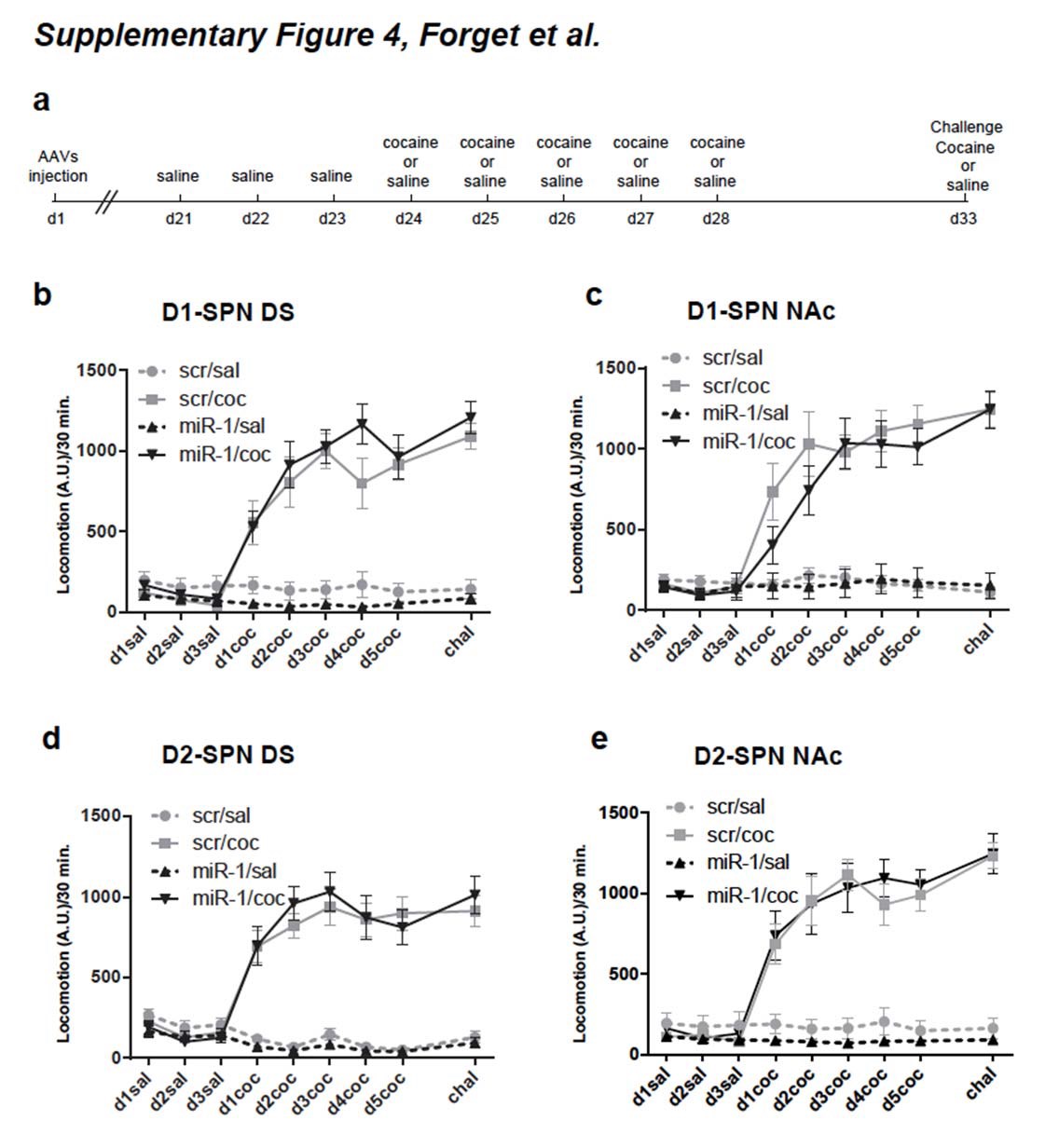
 **Supplementary Figure 4: cocaine-induced locomotor sensitization after miR-1 overexpression in D1- or D2-SPN of the DS or Nac or mice. a)** Schematic representation of the timing of cocaine or saline administrations. **b)** cocaine-induced locomotor sensitization after miR-1 overexpression in D1-SPN of the DS. The 3-ways ANOVA performed on the locomotor activity indicated a significant effect of cocaine treatment (F_1,32_=116.07, p< 0.0001) and cocaine session (F_5,32_=3.27, p=0.0078) but not of miR-1-overexpression (F_1,32_= 0.05, p=0.94); and no interaction (F_1,32_= 0.28, p=0.6). scr/coc: n=10; scr/sal: n=8; miR-1/coc: n=11; miR-1/sal: n=7. **c)** cocaine-induced locomotor sensitization after miR-1 overexpression in D1-SPN of the NAc. The 3-ways ANOVA performed on the locomotor activity indicated a significant effect of cocaine treatment (F_1,36_=55.76, p<0.0001) and cocaine session (F_5,36_=7. 6, p<0.0001) but not of miR-1-overexpression (F_1,36_= 0.38, p=0.54); and no interaction (F_1,36_= 0.34, p=0.56). scr/coc: n=12; scr/sal: n=8; miR-1/coc: n=13; miR-1/sal: n=7. **d)** cocaine-induced locomotor sensitization after miR-1 over-expression in D2-SPN of the DS. The 3-ways ANOVA performed on the locomotor activity indicated a significant effect of cocaine treatment (F_1,30_=83.74, p< 0.0001) and cocaine session (F_5,30_=6.73, p<0.0001) but not of miR-1-overexpression (F_1,30_= 0.004, p=0.95); and no interaction (F_1,30_= 1.29, p=0.26). scr/coc: n=11; scr/sal: n=6; miR-1/coc: n=10; miR-1/sal: n=9. **e)** cocaineinduced locomotor sensitization after miR-1 over-expression in D2-SPN of the NAc. The 3ways ANOVA performed on the locomotor activity indicated a significant effect of cocaine treatment (F_1,40_=65.4, p<0.0001) and cocaine session (F_5,40_=3.2, p=0.0082) but not of miR-1overexpression (F_1,40_= 0.066, p=0.79); and no interaction (F_1,40_= 0.32, p=0.58). scr/coc: n=16; scr/sal: n=5; miR-1/coc: n=14; miR-1/sal: n=7.

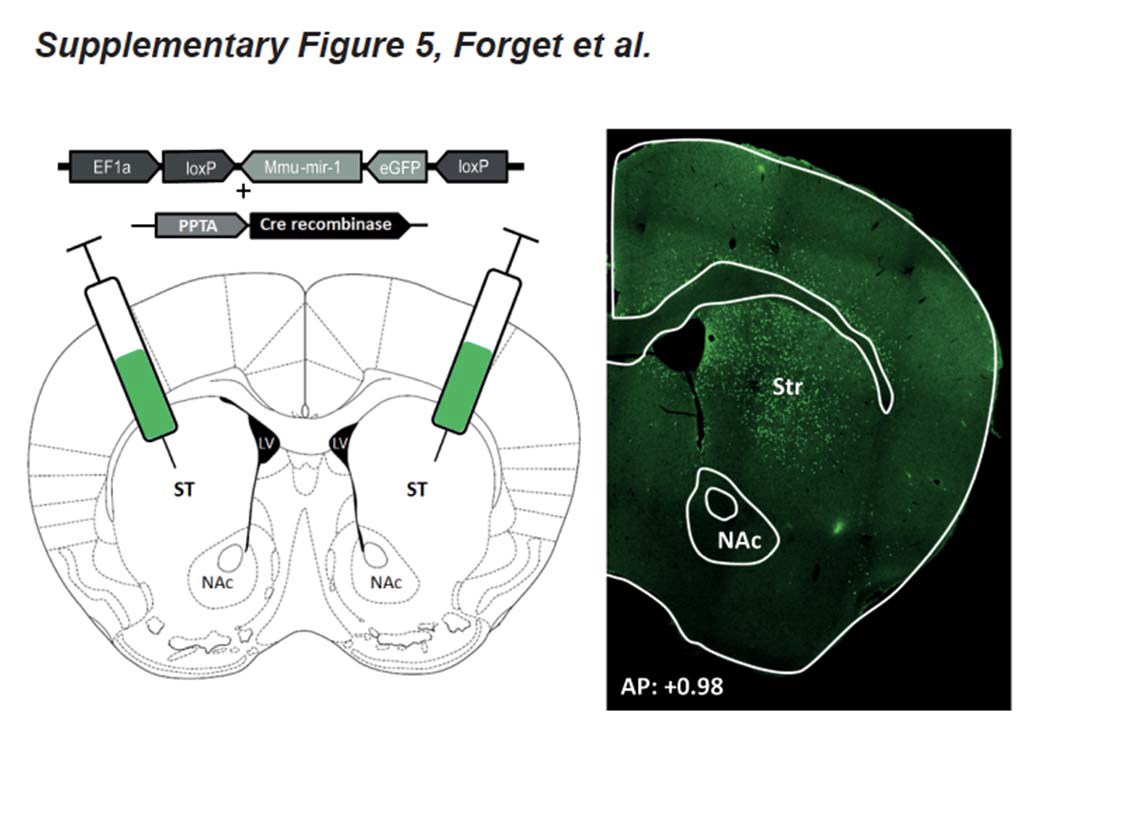
 **Supplementary Figure 5 :** Schematic representation of AAVs injection for miR-1 overexpression in the DS (left panel) and representative image of a brain slice after immunostaining for Cre-recombinase (green, right panel) 24h after the test for cue-induced reinstatement of cocaine seeking (self-administration experiment).

| **Figures** | **Statistical test** | **results** |
| --- | --- | --- |
| Fig. 1b | Mann-Whitney | Left (n=4 per group): all miRNAs: U=8, p>0.99; except miR-124 (U=7, p=0.77).  Right (n=5 per group): drosha (U=10, p=0.6); dicer (U=5, p=0.12); trbp (U=9, p=0.46); ago2 (U=8, p=0.35) |
| Fig. 1c | Mann-Whitney | Left (n=5 per group): all miRNAs: U=0, p=0.14; except miR-124 (U=4, p=0.076), miR-128b (U=2, p=0.05) and let-7d (U=10, p=0.6).  Right (n=5 per group): drosha (U=8.5, p=0.71); dicer (U=9, p=0.8); trbp (U=6, p=0.33); ago2 (U=7, p=0.46). |
| Fig. 1e | Mann-Whitney | DS (n=5 per group): bdnf (U=8, p=0.35); fosb (U=1, p=0.016); npas4 (U=2, p=0.05); gria2 (U=9, p=0.46); grin2b (U=7, p=0.25); gria3 (U=7, p=0.25); pi3k (U=0, p=0.009); D1R (U=10, p=0.6); mef2c (U=7, p=0.46).  NAc (n=5 per group): bdnf (U=2, p=0.05); fosb (U=4, p=0.14); npas4 (U=4, p=0.14); gria2 (U=4, p=0.14); grin2b (U=5, p=0.22); gria3 (U=6, p=0.32); pi3k (U=6, p=0.32); D1R (U=7, p=0.46); mef2c (U=9, p=0.8). |
| Fig. 2a | Mann-Whitney | Bdnf (n=4 per group; U=0, p=0.02); fosb (n=5 per group; U=0, p=0.009); npas4 (n=7 per group; U=0, p=0.0027) |
| Fig. 2b | Mann-Whitney | DS: mir-1 (n=11) *vs* mir-scr (n=12). miR-1 level: U=0, p<0.0001. bdnf: U=42.5, p=0.15; fosb: U=11, p=0.0007; npas4: U=12, p=0.0009.  NAc: n=5 per group. miR-1 level: U=0, p<0.0079. bdnf: U=6, p=0.17; fosb: U=0, p=0.009; npas4: U=0, p=0.009. |
| Fig. 2c | Mann-Whitney | DS: n=5 per group. miR-1 level: U=0, p<0.0079. bdnf: U=6.5, p=0.39; fosb: U=3, p=0.086; npas4: U=1, p=0.027.  NAc: miR-1 level (n=5 per group): U=0, p<0.0079. mir-1 (n=4) *vs* mir-scr (n=5): bdnf: U=5, p=0.22; fosb: U=3, p=0.086; npas4: U=2, p=0.05. |
| Fig. 3a | Kolmogorov Smirnov (KS) normality test | CPP index: p=0.2 for all groups, NS.  Extinction: coc/scr (p=0.2, NS); coc/miR-1 (p=0.16, NS)  Reinstatement index: sal/scr (p=0.2, NS) ; sal/miR-1 (p=0.2, NS) ; coc/scr (p=0.076, NS); coc/miR-1 (p=0.2, NS) |
| Fig. 3a | Two-way ANOVAs | CPP index: treatment (F_1,63_ = 14.9, p = 0.0003), AAV (F_1,63_ = 0.16, p=0.69, NS), interaction (F_1,63_ = 0.34, p=0.56, NS).  Extinction: session (F_5,200_=5.43, p=0.0001), AAV (F_5,40_=0.07, p=0.79, NS), interaction (F_5,200_=0.41, p=0.84, NS).  Reinstatement index: treatment (F_1,63_ = 6, p = 0.017), AAV (F_1,63_ = 1.57, p=0.21, NS), interaction (F_1,63_ = 1.82, p=0.18, NS). |
| Fig. 3a | Bonferroni’s multiple comparison test | CPP index: sal/scr (n=13) *v*s coc/scr groups (n=22): p=0.002. sal/miR-1 (n=12) *vs* coc/miR-1 (n=20) : p=0.026.  Reinstatement index: sal/scr (n=13) *v*s coc/scr groups (n=22): p=0.0078. sal/miR-1 (n=12) *vs* coc/miR-1 (n=20) : p=0.45, NS. |
| Fig. 3b | KS normality test | miR-1 level: sal/scr (p=0.04), sal/miR-1 (p=0.0004), coc/scr (p=0.0024), coc/miR-1 (p=0.0082). |
| Fig. 3b | Kruskal-Wallis | miR-1 level: sal/scr (n=13), sal/miR-1 (n=12), coc/scr (n=22), coc/miR-1 (n=20).  Group effect: p<0.0001 |
| Fig. 3b  miR-1 levels | Dunn’s multiple comparison test | sal/scr *vs* coc/scr: p>0.99, NS;  coc/scr *vs* coc/miR-1 : p<0.0001;  sal/scr vs sal/miR-1: p=0.0007. |
| Fig. 3b  target levels | Kruskal-Wallis | sal/scr (n=11), sal/miR-1 (n=4), coc/scr (n=10), coc/miR-1 (n=10).  Npas4: group effect: p=0.001.  Fosb: group effect: p=0.0063. |
| Fig. 3b  target levels | Dunn’s multiple comparison test | sal/scr *vs* coc/scr : npas4: p=0.0147; fosb: p=0.028.  coc/scr *vs* coc/miR-1 : npas4: p=0.0026; fosb: p=0.012. |
| Fig. 3c | KS normality test | CPP index: p=0.2 for all groups, NS. Extinction: p=0.2 for all groups, NS.  Reinstatement index: sal/scr (p=0.16, NS); sal/miR-1 (p=0.2, NS); coc/scr (p=0.2, NS); coc/miR-1 (p=0.2, NS). |
| Fig. 3c | Two-way ANOVAs | CPP index: treatment (F_1,39_ = 10.35, p = 0.0026), AAV (F_1,39_ = 0.15, p=0.69, NS), interaction (F_1,39_ = 0.001, p=0.97, NS).  Extinction: session (F_5,110_=0.8, NS), AAV (F_1,22_=0.12, p=0.73, NS), interaction (F_5,110_=0.67, p=0.65, NS).  Reinstatement index: treatment (F_1,39_ = 10.41, p = 0.0025), AAV (F_1,39_ = 0.084, p=0.77, NS), interaction (F_1,39_ = 0.066, p=0.8, NS). |
| Fig. 3c | Bonferroni’s multiple comparison tests | CPP index: sal/scr (n=10) *v*s coc/scr groups (n=11): p=0.027. sal/miR-1 (n=9) *vs* coc/miR-1 (n=13) : p=0.03.  Reinstatement index: sal/scr (n=10) *v*s coc/scr groups (n=11): p=0.019. sal/miR-1 (n=9) *vs* coc/miR-1 (n=13) : p=0.04. |
| Fig. 4a | KS normality tests | CPP index: p=0.2 for all groups, NS.  Extinction: p=0.2 for all groups, NS.  Reinstatement index: sal/scr (p=0.2, NS) ; sal/miR-1 (p=0.2, NS) ; coc/scr (p=0.16, NS); coc/miR-1 (p=0.2, NS). |
| Fig. 4a | Two-way ANOVAs | CPP index: treatment (F_1,36_ = 9.92, p = 0.0033), AAV (F_1,36_ = 0.035, p=0.85, NS), interaction (F_1,36_ = 0.16, p=0.39, NS).  Extinction: session (F_5,95_=2.63, p=0.028), AAV (F_1,19_=0.029, p=0.86, NS), interaction (F_5,95_=2.07, p=0.075, NS).  Reinstatement index: treatment (F_1,36_ = 15.48, p = 0.0004), AAV (F_1,36_ = 0.86, p=0.36, NS), interaction (F_1,36_ = 0.72, p=0.4, NS). |
| Fig. 4a | Bonferroni’s multiple comparison tests | CPP index: sal/scr (n=10) *v*s coc/scr groups (n=11): p=0.048. sal/miR-1 (n=9) *vs* coc/miR-1 (n=13) : p=0.022.  Reinstatement index: sal/scr (n=10) *v*s coc/scr groups (n=11): p=0.028. sal/miR-1 (n=9) *vs* coc/miR-1 (n=13) : p=0.003. |
| Fig. 4b | KS normality tests | CPP index: p=0.2 for all groups, NS.  Extinction: p=0.2 for coc/scr, NS; p=0.16 for coc/miR-1, NS.  Reinstatement index: sal/scr (p=0.2, NS) ; sal/miR-1 (p=0.2, NS) ; coc/scr (p=0.085, NS); coc/miR-1 (p=0.2, NS). |
| Fig. 4b | Two-way ANOVAs | CPP index: treatment (F_1,33_ = 27.8, p<0.0001), AAV (F_1,33_ = 0.17, p=0.68, NS), interaction (F_1,33_ = 0.52, p=0.48, NS).  Extinction: session (F_5,90_=4.92, p=0.0005), AAV (F_1,18_=0.05, p=0.83, NS), interaction (F_5,90_=0.7, p=0.62, NS).  Reinstatement index: treatment (F_1,33_ = 11.658, p = 0.0017), AAV (F_1,33_ = 0.94, p=0.34, NS), interaction (F_1,33_ = 0.03, p=0.86, NS). |
| Fig. 4b | Bonferroni’s multiple comparison tests | CPP index: sal/scr (n=9) *v*s coc/scr groups (n=10): p=0.0025. sal/miR-1 (n=8) *vs* coc/miR-1 (n=10) : p=0.0002.  Reinstatement index: sal/scr (n=9) *v*s coc/scr groups (n=10): p=0.026. sal/miR-1 (n=8) *vs* coc/miR-1 (n=10) : p=0.018. |
| Fig. 5a | KS normality tests | Active nose spokes: p=0.13 for control (n=11), NS; p=0.2 for miR-1 (n=11), NS.  Infusions: p=0.2 for both groups, NS. |
| Fig. 5a | Two-way ANOVAs | Active nose spokes: session (F_9,180_ = 37.9, p<0.0001), AAV (F_1,20_ = 0.81, p=0.38, NS), interaction (F_9,180_ = 0.72, p=0.69, NS).  Infusions: session (F_9,180_=5.47, p<0.0001), AAV (F_1,20_=0.72, p=0.4, NS), interaction (F_9,180_=1.01, p=0.43, NS). |
| Fig. 5b | Mann-Whitney | Breaking point: n=11 per group. Control *vs* miR-1: U=41.5, p=0.19, NS. |
| Fig. 5c | KS normality test | Extinction: every session passed the normality test. |
| Fig. 5c | Two-way ANOVAs | Extinction: session (F_19,380_ = 24.5, p<0.0001), AAV (F_1,20_ = 1.4, p=0.25, NS), interaction (F_19,380_ = 0.86, p=0.63, NS). |
| Fig. 5d | KS normality test | Extinction: p=0.2, NS.  Cue-reinstatement: p=0.2, NS. |
| Fig. 5d | Two-way ANOVAs | Cue reinstatement (F_1,20_ = 84.3, p<0.0001), AAV (F_1,20_ = 5.62, p=0.028), interaction (F_1,20_ = 4.78, p=0.04. |
| Fig. 5d | Bonferroni’s multiple comparison tests | Extinction vs cue: reinstatement: p=0.0003 (scr, n=11); p<0.0001 (miR-1, n=11)  Control vs miR-1: p=0.17 (extinction, NS); p=0.033 (cue-reinstatement). |
| Fig. 5e | Mann-Whitney | Scr (n=5) vs miR-1 (n=6): StrM (U=6, p=0.01, NS); StrL (U=3, p=0.028); Shell (U=5, p=0.068, NS); Core (U=4, p=0.045); AC (U=7, p=0.14, NS). |
| Fig. 5f | KS normality tests | Active nose spokes: p=0.12 for control (n=11), NS; p=0.13 for miR-1 (n=14), NS.  Infusions: p=0.17 for control, p=0.1 for miR-1, NS. |
| Fig. 5f | Two-way ANOVAs | Active nose spokes: session (F_9,207_ = 58.1, p<0.0001), AAV (F_1,23_ = 0.06, p=0.8, NS), interaction (F_9,207_ = 0.5, p=0.87, NS).  Infusions: session (F_9,207_=21.15, p<0.0001), AAV (F_1,23_=0.65, p=0.43, NS), interaction (F_9,207_=0.24, p=0.99, NS). |
| Fig. 5g | Mann-Whitney | Breaking point: n=11 for Control, n=14 for miR-1; Control *vs* miR-1: U=37.5, p=0.018. |
| Fig. 5h | KS normality test | Extinction: every session passed the normality test. |
| Fig. 5h | Two-way ANOVAs | Extinction: session (F_19,437_ = 20.7, p<0.0001), AAV (F_1,23_ = 0.45, p=0.51, NS), interaction (F_19,437_ = 0.69, p=0.83, NS). |
| Fig. 5i | KS normality test | Extinction: p=0.4, NS Cue-reinstatement: p=0.7, NS. |
| Fig. 5i | Two-way ANOVAs | Cue reinstatement (F_1,23_ = 132.8, p<0.0001), AAV (F_1,23_ = 0.25, p=0.62, NS), interaction (F_1,23_ = 0.7, p=0.4, NS). |
| Fig. 5i | Bonferroni’s multiple comparison tests | Extinction vs cue-reinstatement: p<0.0001 (scr, n=11); p<0.0001 (miR-1, n=14)  Control vs miR-1: p>0.99 (extinction, NS); p=0.71 (cue-reinstatement, NS). |

**Table 1 : summary of statistical tests in each principal figures.**
