## supplementary methods for "Cell type- and region-specific modulation of cocaine seeking by micro-RNA-1 in striatal projection neurons"

@Supplementary **MATERIALS AND METHODS**

**Animals and drug treatments**

For the self-administration study, mice were tested during the dark phase of a reverse light cycle (lights off at 8.00 a.m and on at 20.00 p.m). All experimental protocols were performed in accordance with the guidelines of the European Communities Council Directive 2010/63/EU and approved by the local ethical committee (Comitè Ètic d'Experimentació Animal-Parc de Recerca Biomèdica de Barcelona, CEEA-PRBB). In agreement, maximal efforts were made to reduce the suffering and the number of mice used.

For the stereotaxic and intravenous catheter implantation surgeries, ketamine hydrochloride (Imalgène; Merial Laboratorios S.A., Barcelona, Spain) and medetomidine hydrochloride (Domtor; Esteve, Barcelona, Spain) were mixed and dissolved in sterile 0.9% physiological saline and administered intraperitoneally in a volume of 10 ml/kg of body weight. Atipamezole hydrochloride (Revertor; Virbac, Barcelona, Spain) and Meloxicam (Metacam; Boehringer Ingelheim, Rhein, Germany) were dissolved in sterile 0.9% physiological saline and administered subcutaneously in an injection volume of 10 ml/kg of body weight.

Gentamicine (Genta-Gobens; Laboratorios Normon, S.A., Madrid, Spain) was dissolved in sterile 0.9% physiological saline and administered intraperitoneally in an injection volume of 10 ml/kg of body weight.

**Stereotaxic injections:**

Mice were anesthetized using ketamine and xylazine (5:1; 0.01 µL/kg). After surgery of mice, anesthesia was reversed by subcutaneous injection of the synthetic α2 adrenergic receptor antagonist, atipamezole (2.5 mg/kg) indicated for the reversal of the sedative and analgesic effects of medetomidine (α2 adrenergic receptor agonist). In addition, mice received an intraperitoneal injection of gentamicine (1 mg/kg) along with subcutaneous administration of the analgesic meloxicam (2 mg/kg). Mice were then placed in a stereotaxic frame (David Kopf, Tujunga, CA) under sterile conditions, with verification that the skull was horizontal. An incision was made over the skull, thus the skull was surgically exposed, the periosteum removed, and burr holes were subsequently drilled with the size of the guide cannula. All the viral injections were made through a bilateral injection cannula (33-gauge internal cannula, Plastics One, UK) connected to a polyethylene tubing (PE-20, Plastics One, UK) attached to a 10 µl microsyringe (Model 1701 N SYR, Cemented NDL, 26 ga, 2 in, pointstyle 3, Hamilton company, NV). The displacement of an air bubble inside the length of the polyethylene tubing that connected the syringe to the injection needle was used to monitor the microinjections. The target volume of 1 µl in the NAc or 2 µl in the dorsal striatum was injected at a constant rate of 0.20 µl/min during 10 min by using a microinfusion pump (Harvard Apparatus, Holliston, MA). After infusion, the injection cannula was left in place for an additional period of 5 min to allow the fluid to diffuse and to prevent reflux, and then it was slowly withdrawn during 5 additional min. Animals were kept on a 37 °C heating pad during the surgery, and until recovery from anesthesia. The verification of viral vector injection site was performed by measurement of miR-1 expression or by immunofluorescence after the self-administration experiment, where the injection sites were verified under a confocal microscope by an experimenter blind to the AAV. Mice with injection location outside of the target area were excluded from the results.

**Quantitative PCR analysis (qPCR) of miRNA and mRNA levels**

The specificity of the reaction was verified by melt curve analysis. Individual data were normalized using a combination of three housekeeping genes (*Ppia*, *HPRT and Rpl38*) for mRNA levels, and using *snoU6* for miRNA levels. The results are reported as fold changes calculated as the ratios of normalized gene expression data for drug-treated groups in comparison to the control group. All mRNA primers were designed using Primer3 software and target specificity was assessed using NCBI Primer-BLAST. miRNA primers were designed according the method described in (1). All primers were purchased from EuroGenTech.

**Luciferase assay to assess the interaction between miR-1 and the 3'UTR of BDNF, FosB and NPAS4 in HEK cells:**

1 x $10^4$ HEK cells per well (96-well plates) have been plated in 50 µl culture medium (DMEM, high glucose, GutaMAX™ 10% FBS, 1% Penicillin-Streptomycin) from a sub-confluent HEK293T cell suspension (cell density < 1.3 x $10^5$ cells/$cm^2$). 3 wells have been plated for each condition to produce triplicates (conditions included a miRNA negative control which does not target the 3'UTRs).

For 1 well, 300 ng of pmirGLO Dual-Luciferase miRNA Target Expression Vectors have been diluted in 25 µl of pure DMEM in combination with miRNA Mimics at a final

concentration of 20 nM with 0.5 µl of Lipofectamine 2000 Transfection Reagent (Life Technologies) diluted in 25 µl of pure DMEM. The transfection complexes have been incubated 20 min at room temperature and added to the well containing plated cells in 50 µl medium (after 8 h in culture).
The transfected cells have been incubated at 37 °C in a 5% CO2 incubator for 48 h without medium change.

After 48h, the volume of the medium left in one well has been measured by pipetting and the appropriate amount of medium has been discarded to reduce the volume to 50 µl from the well.

50 µl of Dual-Glo® Luciferase Reagent (Promega Corporation) has been added to each well (equal volumes). The Incubation lasted 10 min in the dark at room temperature. The firefly luciferase activity was then measured using the luminometer (FLUOstar Omega Microplate Reader - BMG LABTECH). 50 µl of Dual-Glo® Stop&Glo® Reagent (Promega Corporation) was then added to each well and the incubation lasted 10 min in a dark room. The Renilla luciferase activity was then measured using the luminometer.

**Cocaine-induced locomotor sensitization:**

The locomotor activity was evaluated using a circular corridor (4.5 cm width, 17 cm external diameter) in a low luminosity environment with four infrared beams placed at every 90° (Imetronic, Pessac, France). Locomotor activity was counted when animals interrupted two adjacent beams and, thus, had traveled 1/4 of the circular corridor. Locomotor activity was measured by 5 min intervals and cumulative counts were taken for data analysis.

**Cocaine-induced Conditioned place preference (CPP) and relapse:**

The CPP was evaluated in a Plexiglas Y-shaped apparatus (Immetronic) located in a soundproof testing room with low luminosity; with an unbiased procedure. Two of the three chambers were used for the conditioning and distinguished by different patterns on floors and walls, separated by a central neutral area. The device is connected to an electronic interface for data collection. Entries and time spent in each chamber were measured, as well as the locomotor activity of the mice during the experiments.

### Self-administration study

#### Operant self-administration apparatus

The self-administration experiments were conducted in mouse operant chambers (Model ENV-307A-CT; Med Associates Inc., Georgia, VT, USA) equipped with two holes, one randomly selected as the active hole and the other as the inactive hole. The chambers were made of aluminum and acrylic, with grid floors and were housed in sound- and light-attenuated boxes equipped with fans to provide ventilation and white noise. Pump noise and stimuli lights (cues), one located inside the active hole and the other above it, were paired contingently with the delivery of the reinforcer. Cocaine was infused via a syringe that was mounted on a microinfusion pump (PHM-100A; Med Associates) and connected, via Tygon tubing (0.96 mm o.d., Portex Fine Bore Polythene Tubing, Portex Ltd, Kent, UK) to a single-channel liquid swivel (375/25, Instech Laboratories, Plymouth Meeting, PA, USA) and to the mouse intravenous catheter. The swivel was mounted on a counter-balanced arm above the operant chamber.

#### Operant self-administration of cocaine

Mice were anesthetized with a ketamine (75 mg/kg)/medetomidine (1 mg/kg) mixture then implanted with indwelling intravenous silastic catheters. After surgery of mice, anesthesia was reversed by subcutaneous injection of the synthetic α2 adrenergic receptor antagonist, atipamezole (2.5 mg/kg) indicated for the reversal of the sedative and analgesic effects of medetomidine (α2 adrenergic receptor agonist). Responses on the inactive hole and all responses elicited during the 10-s timeout period were also recorded. Cocaine was infused in 23.5 μl over 2 s (0.5 mg/kg per injection, intravenously). The criteria for self-administration behavior were achieved when all of the following conditions were met: (1) mice maintained stable, responding with 20% deviation from the mean of the total number of reinforcers earned in three consecutive sessions (80% of stability); (2) at least 75% of mice responding on the active hole; and (3) a minimum of 10 reinforcers per session. The criterion for extinction was achieved when, during 3 consecutive sessions, mice completed a mean number of nose pokes in the active hole consisting of 30% of the mean responses obtained during the 3-day period to achieve the acquisition criteria for cocaine self-administration training. The reinstatement criterion was achieved when nose pokes in the active hole were double the number of nose pokes in the active hole during the 3 consecutive days when the mice acquired the extinction criteria.

**Tissue preparation for immunofluorescence**

Mice were deeply anesthetized by i.p. injection (0.2 ml/10 g of body weight) of mixture of ketamine/medetomidine prior to intracardiac perfusion with 4% paraformaldehyde (PFA) in 0.1 M Na2HPO4/ 0.1 M NaH2PO4 buffer (PB), pH 7.5, delivered with a peristaltic pump at 30 ml per min for 2 min. Subsequently, brains were extracted and post-fixed with 4% PFA for 24 h and transferred to a solution of 30% sucrose at 4 °C. Coronal frozen sections (30 µm) of the whole dorsal striatum were obtained on a freezing microtome and stored in a 5% sucrose solution at 4°C until use.

**Immunofluorescence and quantification of Fos-B expression**

Mice were deeply anesthetized by i.p. injection (0.2 ml/10 g of body weight) of mixture of ketamine/medetomidine prior to intracardiac perfusion with 4% paraformaldehyde (PFA). Subsequently, brains were extracted and post-fixed with 4% PFA for 24 h and transferred to a solution of 30% sucrose at 4 °C. Coronal frozen sections (30 µm) of the whole dorsal striatum were obtained on a freezing microtome and stored in a 5% sucrose solution at 4°C until use.
Free-floating slices were rinsed in 0.1 M PB, blocked in a solution containing 3% normal goat serum and 0.3% Triton X-100 in 0.1M PB (NGS-T-PB) at room temperature for 2 h, and incubated overnight at 4ºC in the same solution with the primary antibody to anti-Cre-recombinase (1:500, mouse, MAB3120, Merck Millipore). The next day, after 3 rinses in 0.1 M PB, sections were incubated with the secondary antibody AlexaFluor-488 donkey anti-mouse (1:500, Life Technologies) at room temperature in NGS-T-PB for 2 h. After incubation, sections were rinsed and mounted immediately after onto glass slides coated with gelatine in Fluoromount mounting medium. For the analyze of Fos-B expression, free-floating brain sections were incubated 20 min at room temperature in a solution of TBS containing 0.3% Triton X-100 and then incubated 1h in a blocking solution containing 5% BSA and 0.3% Triton X-100. Brain sections were then incubated overnight at 4°C with a rabbit anti-FosB antibody (Invitrogen Ma5-15056) at 1:500 dilution in TBS. The sections were incubated 2 h at room temperature with Alexa fluor 594-conjugated anti-rabbit secondary antibody (Jackson Immunoresearch) at 1:500 dilution and mounted using Prolong® Gold (Thermofisher scientific).

The images for Fos-b expression were acquired using a using Macro-ApoTome (Zeiss) with a 80x objective. Images were saved with Zen software and the number of Fos-B positive nuclei was automatically counted with the spot detector module of ICY software (Institut Pasteur, Paris, France). This program makes possible to select and count cells automatically without experimenter bias (counts were conducted without knowledge of the group assignments). In each coronal section, we sampled both ipsi- and contralateral regions separately and then averaged the density of the spots for each structure per animal. The results were expressed as number of Fos-B positive cell per square millimeter of cerebral tissue.

### Image analysis

The sections of the brain with Cre-recombinase staining were analyzed with Leica TCS SP5 CFS (fixed stage) upright confocal microscope with two non-descanned HyD detectors. The images of Cre-recombinase staining were processed using the ImageJ analysis software.
